# Assessing methylation detection for primary human tissue using Nanopore sequencing

**DOI:** 10.1101/2024.02.29.581569

**Authors:** Rylee Genner, Stuart Akeson, Melissa Meredith, Pilar Alvarez Jerez, Laksh Malik, Breeana Baker, Abigail Miano-Burkhardt, CARD-long-read Team, Benedict Paten, Kimberley J Billingsley, Cornelis Blauwendraat, Miten Jain

## Abstract

DNA methylation most commonly occurs as 5-methylcytosine (5-mC) in the human genome and has been associated with human diseases. Recent developments in single-molecule sequencing technologies (Oxford Nanopore Technologies (ONT) and Pacific Biosciences) have enabled readouts of long, native DNA molecules, including cytosine methylation. ONT recently upgraded their Nanopore sequencing chemistry and kits from R9 to the R10 version, which yielded increased accuracy and sequencing throughput. However the effects on methylation detection have not yet been documented.

Here we performed a series of computational analyses to characterize differences in Nanopore-based 5mC detection between the ONT R9 and R10 chemistries. We compared 5mC calls in R9 and R10 for three human genome datasets: a cell line, a frontal cortex brain sample, and a blood sample. We performed an in-depth analysis on CpG islands and homopolymer regions, and documented high concordance for methylation detection among sequencing technologies. The strongest correlation was observed between Nanopore R10 and Illumina bisulfite technologies for cell line-derived datasets. Subtle differences in methylation datasets between technologies can impact analysis tools such as differential methylation calling software. Our findings show that comparisons can be drawn between methylation data from different Nanopore chemistries using guided hypotheses. This work will facilitate comparison among Nanopore data cohorts derived using different chemistries from large scale sequencing efforts, such as the NIH CARD Long Read Initiative.

## Introduction

DNA methylation is an epigenetic mechanism that most commonly involves the addition of a methyl group to the fifth carbon of a cytosine residue to form a 5mC (5-methylcytosine) complex. This reversible reaction is catalyzed by DNA methyltransferases (DNMTs) and is positively correlated with the recruitment of histone proteins that compact the DNA structure, making it inaccessible to the transcription machinery. As a result, transcription levels can undergo repression leading to a silencing effect of the associated gene. The dominant form of DNA methylation in terminally differentiated human cells occurs proximal to a guanosine residue, and the resulting loci are often referred to as CpG sites. These CpG sites cluster within promoters to form high methylation frequency regions known as CpG islands (Bird 1986). Methylation differences can impact transcription levels. Methylation patterns can also be inherited and play an important role in cellular processes such as embryo development (Monk et al. 1987; Li et al. 2018), genomic imprinting (Suzuki et al. 2007; Li et al. 1993; Court et al. 2014), X-chromosome inactivation (Sharp et al. 2011), and transcription repression(Moore et al. 2012). As a result, variations in DNA methylation have been associated with human diseases such as aging, neurodegeneration, and cancer (Maschietto et al. 2017; Lunnon et al. 2014; Gasparoni et al. 2018; Smith et al. 2018, 2019; Altuna et al. 2019; Lardenoije et al. 2019; Semick et al. 2019; Wei et al. 2020).

DNA methylation has traditionally been measured using bisulfite conversion(Frommer et al. 1992) followed by methylation arrays or short-read sequencing technologies(Lister et al. 2009). However, these methods have had some limitations in that they: i) can only detect methylation patterns in short genomic intervals/specific genomic regions; ii) are not optimized for identifying methylation in traditionally challenging genomic regions; iii) do not allow for haplotype specific methylation detection; and iv) involve a bisulfite chemical conversion step that can cause DNA degradation(Logsdon et al. 2020). In contrast, long read sequencing technologies developed by Oxford Nanopore Technologies (ONT) and Pacific Biosciences (PacBio) can sequence long native DNA strands ranging from 10-20 kb (both ONT and PacBio) to 1 Mb+ (ONT) in length (Payne et al. 2019). These long reads help improve contiguity and accuracy of genome assembly and they are essential for resolving the traditionally challenging, complex, repetitive regions of the genome. Long-read sequencing has been the driving technology behind advances like the successful sequencing of a complete human centromere (Jain et al. 2018), a T2T human chromosome (Miga et al. 2020; Logsdon et al. 2021), and the first complete human genome (Nurk et al. 2022). Long read sequencing technologies can also directly and simultaneously detect nucleotide modifications such as 5-mC, 5-hmC, and 6-mA without any chemical conversion steps or additional sample processing requirements. These advances open up additional avenues to explore genetic and epigenetic variation through haplotype-specific phasing and allele-specific methylation detection.

The quality and throughput of ONT long read sequencing has rapidly improved over the past decade. The median alignment identity has increased from 85% in 2014 (R9 ONT and chemistry) to 99.7%+ in 2023 (R10 Nanopore Duplex chemistry). Throughput has also increased to the point where sequencing a 30x+ genome is routinely achievable using a single PromethION flow cell (Kolmogorov et al. 2023). While these improvements present promising opportunities for future genomics studies, they pose certain challenges for ongoing, multi-year sequencing projects.The NIH Intramural Center for Alzheimer’s and Related Dementias (CARD) is in the process of sequencing thousands of brain tissue samples from donors with and without Alzheimer’s disease. The first sample cohort was sequenced in 2023 using ONT R9 chemistry and kits. However all subsequent samples are being sequenced with updated R10 chemistry and kits.

The combination of a new nanopore flow cell configuration, updated chemistry, and improved basecalling software algorithms have yielded key improvements in accuracy, especially in homopolymer-rich regions (Sereika et al. 2022; Kolmogorov et al. 2023). This resulted in documentable improvements in resolving haplotypes, structural variants, and SNVs. Additionally, a custom R10 pore-specific methylation calling model was released. ONT stated that the training for the methylation detection model, Remora, that identifies changes in ionic current created by modified and unmodified bases moving through the pore was more comprehensive and robust with R10 chemistry (ONT website). While some of the improvements between the R9 and R10 chemistry-derived data have been documented (Kolmogorov et al. 2023), differences in methylation detection have not been thoroughly documented by the academic community.

In this work, we computationally evaluated methylation calls for a cell line, a brain sample, and a blood sample that were sequenced using both R9 and R10 ONT chemistries. We also used PacBio and Illumina bisulfite sequencing data for the cell line to benchmark methylation calls using orthogonal data. A large motivation behind this work is that some of the CARD samples have been sequenced with R9 and others with R10. Given that many of these samples are limited in quantity, it is important to validate how differences in chemistry and sequencing modalities could affect methylation calling and downstream analyses. Such validations could be used for assessing and implementing strategies for switching over to newer chemistries as technology improves. It can also help devise analysis strategies for comparing variant calling and methylation calling information from different ONT chemistries.

## Results

### ONT sequencing improvements with R10

We first compared sequencing data for an established Genome in a Bottle (GIAB) human cell line (HG002). In our recent work, we sequenced DNA from this cell line using a protocol optimized for the R9.4.1 chemistry and analyzed the data using a custom analysis pipeline for ONT data called NAPU(Kolmogorov et al. 2023). These data had 42x genome coverage and 28 kb read N50. We then sequenced DNA from the same cell line using the R10.4.1 chemistry. The resulting data had 45x genome coverage and 29 kb read N50. We performed benchmarking analyses using these cell line-derived data. Additionally, we sequenced two primary tissue samples (a brain and a blood sample) using both R9 and R10 chemistries to further assess performance.

We basecalled these data using Guppy v6.3.8 and then aligned them relative to the GRCh38 reference genome using minimap2 (see Methods). The median alignment identity was 95.05% for R9 and 98.72% for R10 (Supplementary Table 1). We then performed variant calling using DeepVariant and Sniffles2. The R10 data we analyzed had demonstrable improvements in variant calling relative to R9 (Supplementary Tables 2 and 3) which was in agreement with what was previously documented (Kolmogorov et al. 2023).

### Comparing methylation calls between R9 and R10 ONT data

We extracted methylation information from the aligned data using modkit (https://github.com/nanoporetech/modkit) with the extended bed file format and collapsed strands (see Methods). This resulted in 99.22% and 99.09% of the ∼29.17 million CpG sites in GRCh38 being represented by R9 and R10 data respectively. We then filtered the data to only include CpG sites that had a minimum coverage of 20x and maximum coverage of 200x. This filtering resulted in 25,937,319 CpG sites (88.92% of sites represented in GRCh38) in R9 data and 27,021,032 sites (92.63% of sites represented in GRCh38) in R10 data (Supplementary Table 4).

We first compared CpG sites that had been detected in HG002 R9, R10, and bisulfite sequencing datasets with at least 20x coverage and at most 200x coverage. We binned the sites based on their bisulfite methylation frequencies (reads with 5mC at the site of interest / total reads at the site of interest) that were classified into a combination of 5% and 10% methylation frequency intervals (see Methods). Methylation proportions at both the low (0-10% for bisulfite) and high (90-100% for bisulfite) end of the spectrum saw a significant shift (R10 < R9 for 0-10% bisulfite, Mann-Whitney p = 0.0, R10 > R9 for 90-100% bisulfite, Mann-Whitney p = 0.0) in distributions between R9 and R10 proportions (Figure 1a).

**Figure 1.**
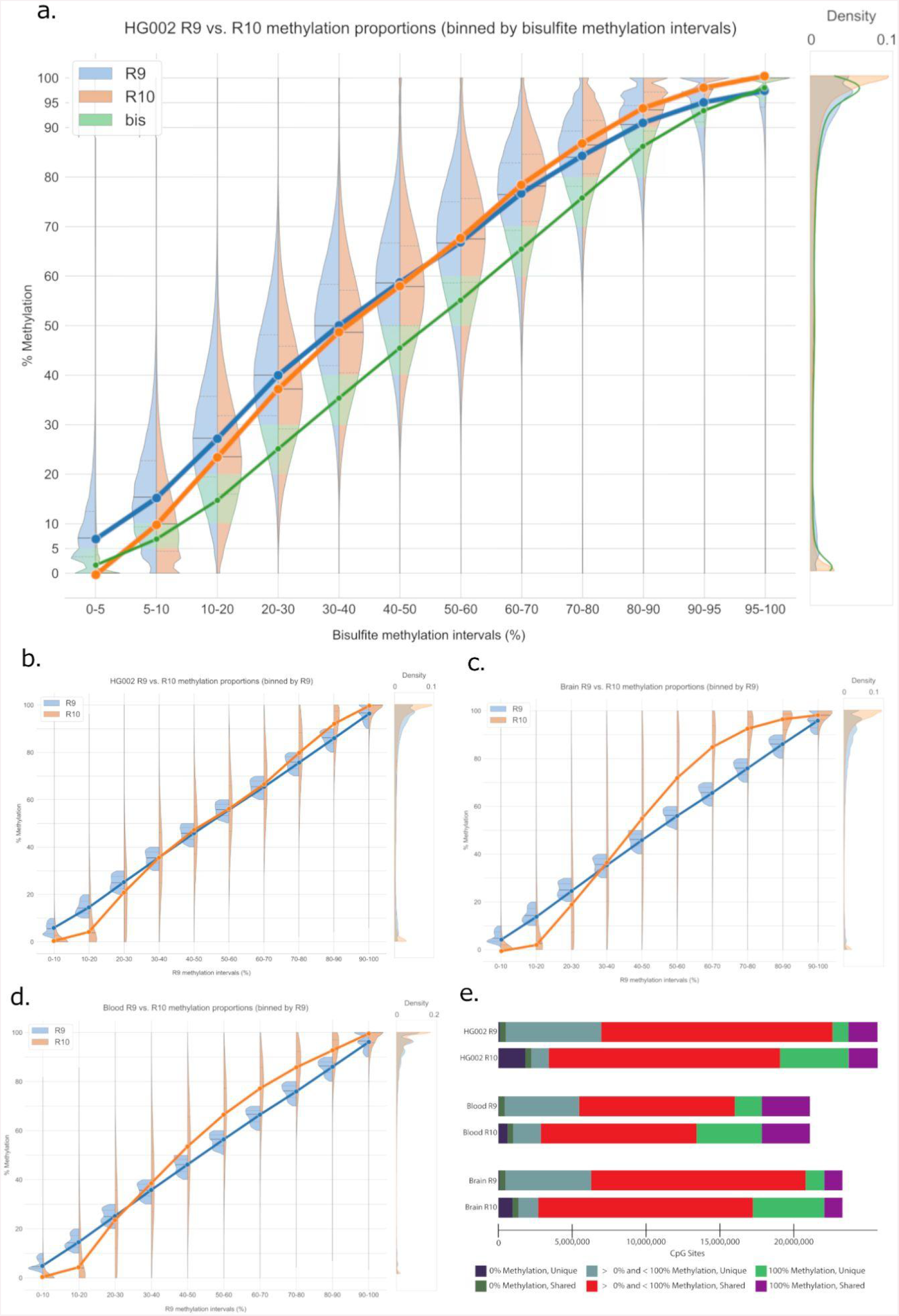
Overall comparison of DNA methylation calls between R9 and R10 datasets across the HG002 cell line, blood, and brain sample. **a**. Methylation proportions of R9 (orange) and R10 (blue) data for the HG002 cell line when binned by bisulfite (green) methylation intervals. The portion of R9 and R10 methylation distributions that agree with the bisulfite methylation range are highlighted in green. The violin plots underneath show methylation proportions of R9 and R10 data for HG002 cell line, brain sample, and blood sample (**b**,**c**,**d**) binned according to R9 intervals. Each violin plot has lines connecting the median interval points for better visualization of methylation trends. Distributions of CpG site methylation frequencies are depicted on the right side of each panel. **e**. Stacked bar charts showing the breakdown of total CpG sites per technology per sample. These sites are further subset into CpG sites with 0% methylation frequency and 100% methylation frequency (Supplementary Table 5-10).

We next compared R9 and R10 performance in the HG002 cell line, brain sample, and blood sample. Each CpG site that we identified in our initial filtering step had a coverage of at least 20 reads and had been detected in both R9 and R10 datasets. We binned the sites based on their R9 methylation proportion and made kernel density estimators to compare distributions between R9 and R10 methylation proportions in each bin. We selected a 0.1 bin size (10% methylation intervals) to minimize the effect of coverage based aliasing on our visual analysis. Methylation proportions at both the low (0-10% for R9) and high (90-100% for R9) end of the spectrum saw a significant (Mann-whitney test p=0.0 for 0.0-0.1 R9 > all R10, Mann-whitney test p=0.0 for 0.9-1.0 R9 < all R10) shift in distributions between the two chemistries (Figure 1 b-d, Supplementary Figure 1 a-c). The two datasets shared 25,521,492 locations after filtering. In R10 data we documented 2,227,118 positions with 0% methylation of which 1,826,337 had > 0% methylation in the R9 data. Similarly of the 6,591,874 positions that had 100% methylation in R10 data, 4,624,404 had < 100% methylation in R9 data. R9 data had fewer positions with 100% or 0% methylation, 3,046,894 and 506,730 respectively. For the 3,046,894 100% methylated positions in R9 data, there were 1,079,424 positions that had < 100% methylation in R10 data. Of the 506,730 positions with 0% methylation for R9 data, 105,949 had > 0% methylation in the R10 data set (Figure 1e, Supplementary Table 5, Supplementary Table 6)

### Characterizing R9 vs. R10 methylation differences

Since R10 ONT data have demonstrated an improved resolution of homopolymers compared to R9 ONT data, we evaluated if this impacted methylation calling. We defined homopolymer regions using the GRCh38 low complexity BED file made available by the Global Alliance for Genomics and Health (GA4GH) Benchmarking Team and the NIST Genome in a Bottle Consortium (([CSL STYLE ERROR: reference with no printed form.]). This bed file defines 3,819,657 homopolymeric regions covering 83,977,437 bases. The intersection between homopolymer regions from HG002 R9 and R10 MethylBed files resulted in 296,660 and 314,448 sites with ≥20x and ≤200x coverage, respectively. Of those sites, 290,592 were shared across both chemistries with a pearson r methylation frequency correlation of 0.966 (RMSE 10.08). Supplementary Figure 2 shows more positions being identified as 0% or 100% methylated in R10 data relative to R9 data, concordant to what we had documented for the whole genome. We also compared methylation frequency in R9 and R10 data for all cytosines genome-wide, documented promoter regions, and CpG islands, and observed a similar pattern (Figure 2 a-c).

**Figure 2.**
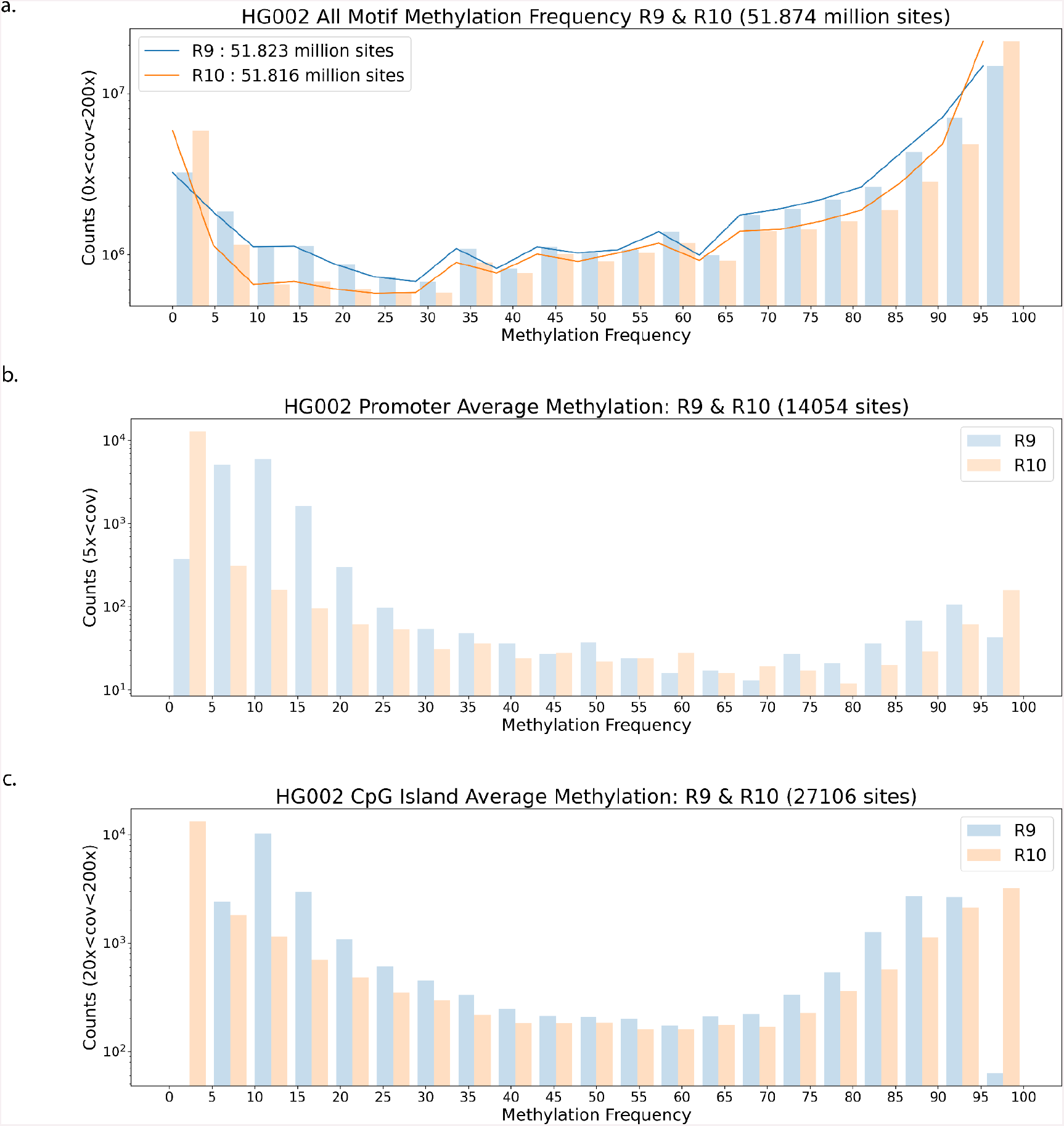
Methylation Calls Compared between R9 and R10 datasets. **a**. Methylation calls for all motifs (all context cytosine) in HG002 R9 and R10, these sites are not strand collapsed. **b**,**c**. panels are average methylation over promoter and CpG island bed regions respectively.

We explored the differences in autosomal protein coding gene promoter regions (Dreos et al. 2017) in HG002 as measured by R9 and R10. The two technologies cover 14,054 of the 29,599 promoters with 5 reads or more with a pearson r methylation frequency correlation of 0.956 (RMSE 10.07) (Figure 2b). Promoters are generally hypomethylated in HG002. R10 reports 9,148 promoters as having 0% methylation while the R9 data peaks with 1,567 promoters having 10% methylation. Overall these regional analyses reveal the same trends as we found genome wide.

To further discern the methylation call differences between R9 and R10, we explored alternative hypotheses. We first examined the impact of strandedness on methylation calling differences (Supplementary Figure 3). We then tested if the shift in proportion we observed was an artifact due to either sample preparation or the sequencing experiment. To that end, we tested an HG002 ultra-long dataset that was sequenced in a different laboratory on a different PromethION device. This comparison, however, yielded the same observation as we had seen from the data generated at CARD (Supplementary Figures 4-8). We also tested if variation in genome coverage between R9 and R10 datasets contributed to methylation calling differences. However, this was not the case (Supplementary Figure 9). We tested then if the difference we observed in methylation proportions could be attributed to the number of CG motifs in a set window size. We did not notice any discrepancies in either a 100 or 1000 base window (Supplementary Figure 10). Finally we examined the covariance of methylation call confidence in reads with at least 20 overlapping CpG sites, showing that R10 had a broader covariance distribution (Supplementary Figure 11).

**Figure 3.**
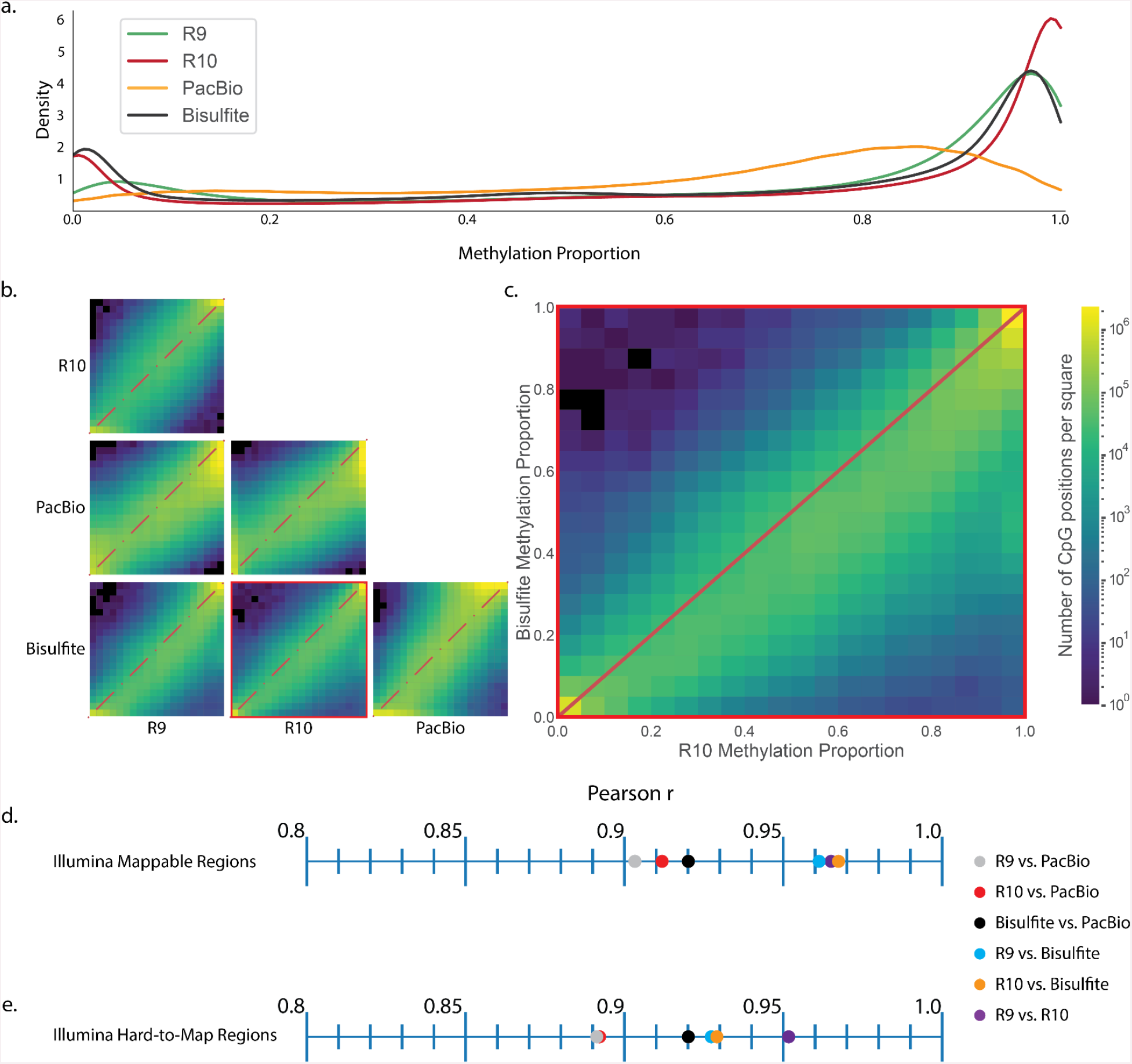
Comparison of HG002 immortalized cell line methylation calling between technologies. **a**. Methylation proportion kernel density estimators for each technology. **b**. Pairwise site specific CpG methylation proportion comparison between technologies. **c**. Site specific CpG methylation proportion comparison between R10 and Bisulfite sequencing. **d**. Pearson r values for pairwise comparisons between technology in Illumina 150 bp paired end mappable regions as defined by BisMap **e**. Pearson r values for pairwise comparisons between technologies in Illumina 150 bp paired end hard-to-map regions.

**Figure 4.**
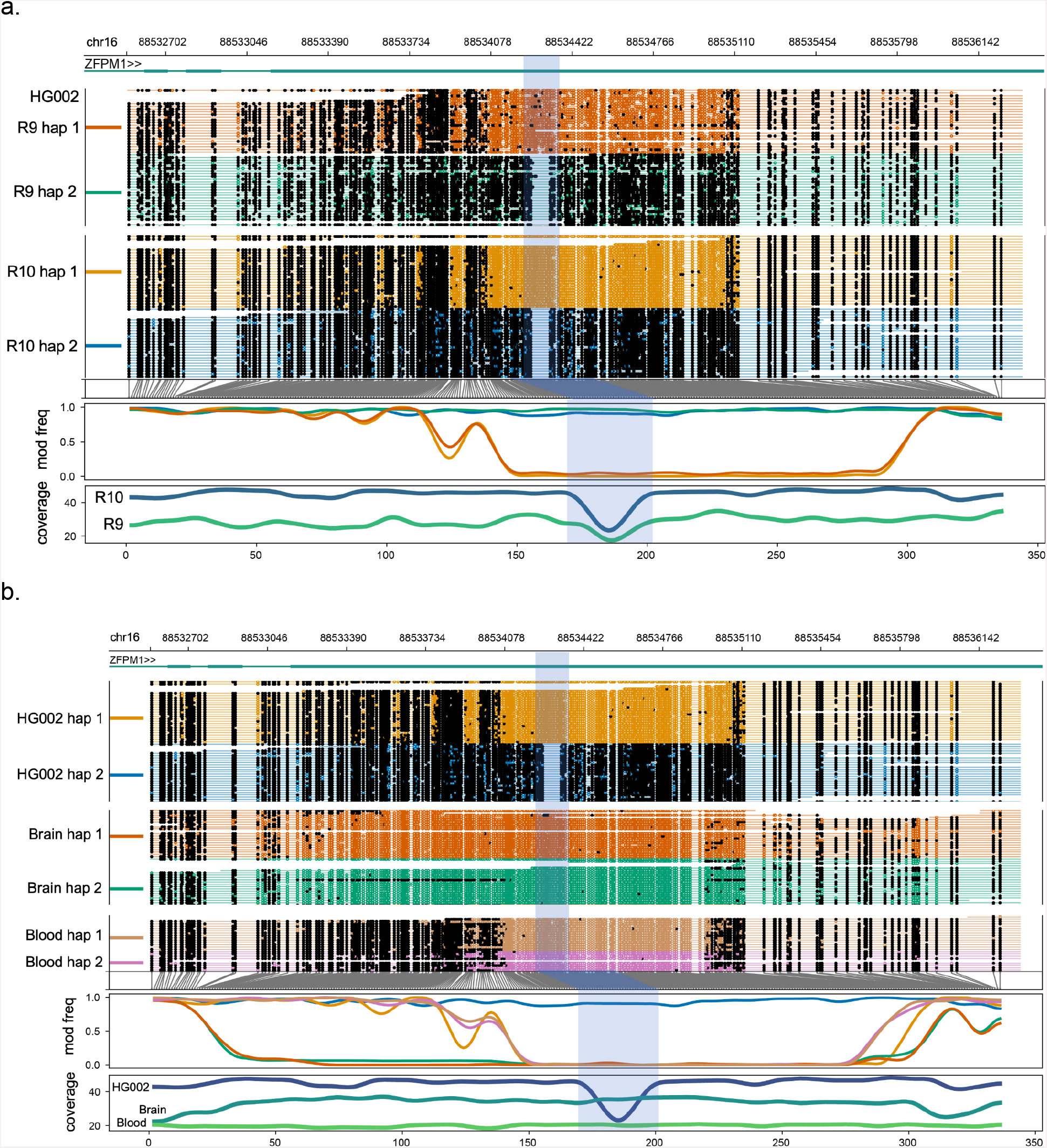
Haplotype-specific methylation differences and similarities between cell, blood and brain samples. From top to bottom, each plot shows the genome coordinates, labeled gene models (if present), haplotype-aware read mappings with modified bases as black (methylated) or colored (unmethylated) circles, a smoothed methylation fraction plot, and a coverage plot. The highlighted region corresponds to a 75 bp deletion (chr16:88534247-88534321) in haplotype 2 of the HG002 cell line that coincides with haplotype-specific methylation. **a**. Haplotype-specific methylation differences and similarities between R9 and R10 sequenced HG002 cell sample. **b**. Haplotype-specific methylation differences and similarities between the R10 sequenced cell, blood and brain samples.

We also compared genome wide CpG site methylation proportions for HG002 datasets generated using ONT’s platform (R9 and R10), PacBio platform (HiFi data), and Illumina platform (bisulfite sequencing data; typically considered to be a “gold standard”) (Figure 3a). We noted that all of the sequencing technologies had good concordance (pearson r >= 0.9) with fluctuations occurring in the methylation proportion of extrema (Figure 3 b-c). Based on these comparisons, ONT R10 and Illumina bisulfite sequencing had the highest overall correlation (r=0.967384 in illumina mappable regions from BisMap), while ONT R9 and PacBio had the lowest correlation (r=0.903295) (Figure 3 d-e). In comparing R9 and R10 we noted that the R9 tended to call the extremes of methylation proportion (0% and 100%) with less frequency than R10. This same phenomenon was observed in varying degrees in each technology we compared to R10.

Additionally, we used Integrative genomics viewer (IGV)(Robinson et al. 2011) to visualize methylation patterns in HG002 cell line between ONT methylation calls (for both R9 and R10 datasets) and traditional bisulfite sequencing in constitutively methylated and constitutively unmethylated regions (Edgar et al. 2014) (Supplementary Figure 12). These examples suggest that the methylation detection differences between chemistries do not hinder their ability to draw qualitative conclusions in a conservative hypothesis context.

We phased ONT reads using PEPPER-Margin-DeepVariant (Shafin et al. 2021) and compared R9 and R10 data for haplotype-specific differential methylation using Nanomethphase(Akbari et al. 2021), which utilizes the R package DSS(Feng et al. 2014). We applied Nanomethphase to calculate differentially methylated regions (DMRs) between R9 and R10 haplotype-phased HG002 samples. R10 identified more DMRs than R9, with similar lengths (Supplementary Table 11-12). The average difference in methylation proportion was higher for R10. This supports our observation that R10 calling the extremes of methylation more frequently than R9 was also contributing to the downstream identification of DMRs.

### Methylation comparison for primary human blood and brain tissue samples

We wanted to assess if ONT data derived from human primary tissue exhibited similar patterns between R9 and R10 chemistries. To that end, we sequenced a human brain sample and a human blood sample using the protocol developed at CARD (see Methods). The data from the brain sample had an average genome coverage of 39x (read N50 30 kb) for R9 and 56x (read N50 26 kb) for R10 chemistry. The data from the blood sample had an average genome coverage of 39x (read N50 34 kb) for R9 and 36x (read N50 37 kb) for R10 chemistry respectively. The median alignment identity for these samples was 95.2% (R9) and 98.52% (R10) for brain-derived and 95.3% (R9) and 98.55% (R10) for blood-derived data (Supplementary Table 1).

For the brain and blood samples we extracted methylation in the same fashion as HG002 (see Methods). For the brain sample this resulted in 98.78% and 98.73% of the ∼29.17 million CpG sites in GRCh38 being represented by R9 and R10 data respectively (Supplementary Table 4). Filtering for 20x coverage or higher resulted in 23,271,407 CpG sites (79.78% of sites represented in GRCh38) in R9 data and 27,742,379 sites (95.11% of sites represented in GRCh38) in R10 data. Of those sites, 23,148,718 overlapped (79.36% of sites represented in GRCh38). For the blood sample we observed 98.12% and 98.07% of the GRCh38 CpG sites for R9 and R10 respectively. After filtering R9 had 22,347,084 CpG sites (76.61% of GRCh38 CpG sites) and R10 had 25,723,371 CpG sites (88.18% of GRCh38 CpG sites). Of those sites 20,977,914 overlapped (71.92% of GRCh38 CpG sites). Additionally we observed a similar pattern of methylation proportions across chemistries in the primary tissue samples (Supplementary Figure 13).

We wanted to assess if there was variation in identification of haplotype-specific methylation between R9 and R10 datasets. This required a strategy to preserve the phasing information between the two datasets because the assignment of haplotype tags is performed at random by PEPPER-MARGIN-Deepvariant. To overcome this limitation, we merged the bam files for R9 and R10 datasets for HG002 cell line and applied PEPPER-MARGIN-Deepvariant (using settings for R9 chemistry) to perform phasing. We then separated the merged and phased R9/R10 haplotagged bam into phased R9 and R10 bam files by filtering for the original R9 and R10 read names. This preserved phase 1 and phase 2 haplotag assignments between the two datasets for downstream comparison. We used modkit to estimate methylation frequencies of the CpG sites and performed differential methylation analysis using the NanoMethPhase dma module. We used the methylartist package to visualize haplotype-specific methylation differences associated with a 75bp deletion on chromosome 16 in the R9 and R10 HG002 cell line datasets (Figure 4a), the R10 HG002 cell line, blood, and brain sample datasets (Figure 4b), and previous R10 HG002, HG02723 and HG00733 GIAB cell line sample datasets (Kolmogorov et al. 2023) (Supplementary Figure 14). We also visualized haplotype-specific methylation differences in the R10 HG002 cell line, blood, and brain samples in the imprinted GNAS region on chromosome 20 (Supplementary Figure 15).

## Discussion

There are several large scale human genome projects underway in the US and across the world. One of them is being led by NIH’s Center for Alzheimer’s Disease and Related Dementias (CARD). As part of this effort, CARD Long Read Initiative, researchers involved with CARD have developed protocols designed to streamline and automate the tissue processing and long-read ONT sequencing of thousands brain samples from individuals with and without Alzheimer’s Disease (AD). These sequencing data provide a unique opportunity to perform genome-wide, population-scale, methylation analyses and assess methylation levels in poorly resolved genomic regions in the human brain. However, as sequencing technologies are continuously improved upon these large scale initiatives may want to use the most up-to-date versions. This will result in cohorts of data sequenced with different sequencing technologies, like R9 and R10 for NIH CARD. It is imperative to document the differences in methylation measurements arising due to technology improvements so that they are not misinterpreted as cohort specific observations.

In this work, we systematically assessed the performance of ONT sequencing for methylation analysis using datasets for cells, blood, and brain tissue from both R9 and R10 chemistries. We also compared ONT methylation detection with other sequencing platforms (ONT, Pacific Biosciences, and Illumina). These comparisons revealed that the overall differences between R9 and R10 methylation datasets were significant enough that they should be taken into account when comparing datasets across platforms and chemistries. Biologically relevant conclusions for methylation across cohorts sequenced using these two chemistries must account for these differences. We argue that long-read sequencing can be an equivalent alternative for methylation to short-read bisulfite sequencing, and include a larger scope of genomic targets.

Direct, simultaneous analysis of modifications will allow for exploring beyond genomic and structural variation in samples across cell types and tissue. This will be transformative for studying biology and the understanding of disease mechanisms. It is also important to characterize differences across different platforms and between technological improvements. Historically, such analyses have focused only on 5mC in CpG contexts. This is especially true of long read technologies. However, ONT sequencing is now capable of detecting 5mC and 5hmC simultaneously. A comprehensive, genome-wide, and context-agnostic analysis of cytosine modifications in human primary tissue samples will be essential for improving our understanding of basic and disease biology. Our analysis strategy can also extend to other modifications as their informatics inference becomes amenable in sequencing data.

## Methods

### Samples collection and sequencing

Long-read sequencing data was generated from human blood, brain and cell-line samples. For the blood sample, frozen blood was obtained from the PPMI study (https://www.ppmi-info.org/) of a 56 year old female donor without known neurological symptoms. For the brain sample, frozen tissue was obtained from the frontal cortex of a 86 year old male donor without known neurological symptoms at the Banner Sun Health Research Institute (https://www.bannerhealth.com/services/research/locations/sun-health-institute/programs/body-donation/tissue). For the cell-line, the HG002 cell-line was purchased from Coriell (https://www.coriell.org/): HG002 (Ashkenazi Jewish ancestry, catalog no. GM24385) and cell culture was performed using Epstein–Barr virus (EBV)-transformed B lymphocyte culture in RPMI-1640 medium with 2 mM l-glutamine and 15% FBS at 37°C.

For DNA processing, the blood(J Billingsley 2022; Miano-Burkhardt 2023), brain(J Billingsley et al. 2022; Baker 2023) and cell line(Alvarez Jerez 2023; Cogan 2023) protocols are explained in detail and are publicly available on protocols.io. In brief, DNA was extracted using either the Nanobind Tissue Big DNA kit (cell line and brain) or the Nanobind HT 1ml blood kit (blood) (PacBio). For the cell line and blood samples, the DNA went through a size selection step using a SRE Kit (PacBio, SKU-102-208-300) to remove fragments up to 25kb. DNA was then sheared to a target size of 30 kb on a Megaruptor3 instrument (Diagenode) with either the DNAFluid+ needles at speed 45 for two cycles (cell and brain) or speed 20 for two cycles with the standard shearing kit (blood). For all samples, DNA length was assessed by running 1 μl on a genomic screentape on the TapeStation 4200 (Agilent). DNA concentration was assessed using the dsDNA BR assay on a Qubit fluorometer (ThermoFisher). Libraries were constructed using either an SQK-LSK 110 kit (ONT) or SQK-LSK 114 kit (ONT) and were loaded onto R.9.4.1 or R.10.4.1 flow-cells respectively. Each sample was sequenced for a total of 72 hours, with roughly one reload every 24 hours on a PromethION device per the manufacturer’s guidelines (ONT, FLO-PRO002).

R9 samples were basecalled using Guppy v6.1.2 (with config file dna_r9.4.1_450bps_modbases_5mc_cg_sup_prom.cfg) and R10 samples were basecalled using Guppy v6.3.8 (with config file dna_r10.4.1_e8.2_400bps_modbases_5mc_cg_sup_prom.cfg). The read batch size and reads per fastq were both set to 50000 and chunks per runner was set to 195 for both R9 and R10.

Example commands below:

R9:

~~~
guppy_basecaller -i ${FAST5_PATH} -s ${OUT_PATH} -c
dna_r9.4.1_450bps_modbases_5mc_cg_sup_prom.cfg -x cuda:all -r
--read_batch_size 50000 -q 50000 --chunks_per_runner 195 --bam_out
~~~

R10:

~~~
guppy_basecaller -i ${FAST5_PATH} -s ${OUT_PATH} -c
dna_r10.4.1_e8.2_400bps_modbases_5mc_cg_sup_prom.cfg -x cuda:all -r
--read_batch_size 50000 -q 50000 --chunks_per_runner 195 --bam_out
~~~

### Structural variant calling

Structural variants were called using sniffles v2.2, a structural variant caller designed for long-read sequencing data. Methylation-tagged bams mapped to GRCh38 were used as the input and the minimum SV length was set to 50 bps.

### CpG site Methylation Frequency Estimation

CpG site methylation frequencies were estimated using modkit, a suite of tools for manipulating ONT modified-base data stored in BAM files. The Modkit pileup command was used with either phased or unphased mapped bams as input to create summary counts of modified and unmodified bases in an extended bedMethyl format - a series of columns detailing the counts of base modifications in each sequencing read over each reference genomic position. Output was restricted to 5mC sites with a CG dinucleotide in the reference and reported. Methylation calls were aggregated/collapsed across strands. (https://github.com/nanoporetech/modkit)

~~~
modkit pileup --cpg --ref --only-tabs --ignore h --combine-strands <IN_BAM>
<OUT_BEDMETHYL>
~~~

### Comparison of R9, R10, Bisulfite and HiFi Methylation proportions genome wide

The unaligned bam files were aligned to the GRCh38 human reference genome([CSL STYLE ERROR: reference with no printed form.]) using a combination of samtools to extract methylation aware fastqs (-TMm,Ml,MM,ML), minimap2 to align fastqs to reference genome (-x map-ont) and samtools again to sort and index aligned bam files. Modkit ([CSL STYLE ERROR: reference with no printed form.]) was used to produce bedMethyl files with collapsed strands from the aligned bam files. A set of Numpy arrays were created and populated with CpG positions from the reference genome, ratios of modified sites calculated as Modified Calls / (Modified Calls + Non-modified Calls), and coverage (Modified Calls + Non-modified Calls).

The split violin plot for Figure 1a was created by filtering the modkit bedMethyl files for CpG sites shared between R9, R10 ONT technologies and a bisulfite bedMethyl file. Only CpG sites on the main chromosomes (1-22, X, Y, M) with coverage levels between 20x and 200x for all three datasets were considered. A pandas dataframe was created with each row featuring a CpG site (defined by genomic coordinates) and its accompanying information. The “pandas.cut” function was used to bin CpG site methylation frequencies into specified intervals ([0-5), [5-10), [10-20), [20-30), [30-40), [40-50), [50-60), [70-80), [90-95), [95-100)) based on the bisulfite dataset, with the rightmost edge values included. Split violin plots were used to plot the methylation frequency distributions across each interval, with the R9 distribution on the left in blue and the R10 distribution on the right in orange. The intervals were classified on the x-axis and the actual distribution of values within those intervals were on the y-axis. A segmented line plot of the median value for each technology at each interval was drawn. A smoothed histogram comparing the distribution of methylation frequencies within each technology (R9 in blue, R10 in orange and bisulfite in green) was added along the right side of the graph (Figure 1a). The additional split violin plots were created in the same manner, but were binned into 10% intervals by R9 instead of bisulfite in Figure 1 b-d and by R10 in Supplementary Figure 1 a-c.)

Pearson r values were calculated for three sets of the data; all CpG sites meeting coverage filtering criteria, all CpG sites meeting coverage filtering criteria in illumina mappable (150bp paired end reads(Hoffman)), and all CpG sites meeting coverage filtering criteria not in the illumina mappable regions.

The heatmaps were created by plotting methylation proportions for each genomic site for one technology on the x-axis and another technology on the y-axis in bins equating to 0.05 sized buckets. Each combination of R9, R10, Bisulfite, and HiFi data were plotted.

This process was repeated for R10 and Pacbio HiFi Methylation data, and R10 and illumina data. This process was repeated with the added caveat of separating CpG sites by strand of origin, creating a paired violin plot for both strands at each of the R10 binned proportions.

### CpG Density and Coverage Analysis

Scatterplots of difference in coverage and difference in proportion of methylation were created using the same strategy as listed above. These arrays were plotted as the x and y-axis respectively, with hue representing the density of observed values.

CpG density plots were created by calculating the number of CpG sites around each CpG site, including the CpG site of origin. This process was repeated in 10, 100, and 1000, nucleotide windows centered at the C of the CpG site of origin. The number of CpG sites in the accordant window was used as the x-axis, while the difference in methylation proportion between R9 and R10 was used as the y-axis.

### Phased Differentially Methylated Regions; R9 vs. R10

For each sample, the R9 and R10 GRCh38-mapped bams were merged and phased together using PEPPER-MARGIN-Deepvariant with R9 settings (-ont_R9_guppy5_sup flag). This was done to keep phase 1 and phase 2 assignments consistent since they are normally randomly assigned by PMVD. The merged and phased combined R9/R10 haplotagged bam was then separated into phased R9 and R10 bam files by filtering for the original R9 and R10 read names using Picard -FilterSamRead (part of the GATK toolbox).

Differentially methylated regions were calculated by using Nanomethphase dma, a python package built on top of the Bioconductr DSS library. Comparisons between phased-haplotypes for the same chemistry (eg. R10 Haplotype 1 vs. R10 Haplotype 2) and between chemistries for the same haplotype (eg. R9 Haplotype 1 vs R10 haplotype 1) were performed for the HG002 cell line data.

Comparisons between DMRs were done with bedtools intersect.

~~~
bedtools intersect -wao -a {file1} -b {file2} > {outfile}
~~~

This produced a bed file from which counts of overlapping bases for each pair of intersecting DMRs between chemistries was calculated. This process needs to be repeated with the inverse orientation of bedfiles since the overlap calculation is not a symmetric function.

### Covariance Calculations

Covariance was calculated by finding all reads with 20 or more CpGs overlapping and inputting paired methylation confidence calls (ML tags in the bam files) into the covariance formula Cov(X,Y)=E[(X−EX)(Y−EY)]=E[XY]−(EX)(EY).

### Haplotype-specific DMR Visualization

Haplotype-specific methylation differences were visualized using Methylartist, a tool for parsing and plotting methylation patterns from ONT data. Mapped bam files were used as input and the locus command was used to generate haplotype-aware, smoothed methylation profiles across specified intervals. A coverage plot was added and the raw log-likelihood ratio section was excluded from the final graph. https://github.com/adamewing/methylartist

~~~
methylartist locus -b <IN_BAM1>,<IN_BAM2> -i chr16:88532537-88536321 -g <REF>
—-plot_coverage <IN_BAM1>,<IN_BAM2> —-labelgenes –-genes ZFPM1 –-motif CG
—-phased –slidingwindowsize 5 –-samplepalette colorblind --nomask
--coverpallete viridis -—ignore_ps -o <OUT_PREFIX>
~~~

## Supporting information

Supplement - R9 and R10 5mC manuscript

## Data availability

### Blood and Brain

Blood and Brain sequencing data is currently in the process of being uploaded to https://anvilproject.org/ accessible via https://terra.bio/ and will be made available as well at https://www.alzheimersdata.org/ad-workbench

Human brain sequencing datasets are under controlled access and require a dbGap application (phs001300.v4). Afterwards, the data will be available through the restricted AnVIL workspace: https://anvil.terra.bio/#workspaces/anvil-datastorage/ANVIL_NIA_CARD_LR_WGS_NABEC_GRU.

### HG002 Bisulfite

HG002 Illumina bisulfite sequencing data were collected from the an AWS open data set generated by ONT s3://ont-open-data/gm24385_mod_2021.09/ described here (https://labs.epi2me.io/gm24385-5mc).

### HG002 PacBio HiFi

HG002 HiFi data are available through Genome in a Bottle described here https://ftp-trace.ncbi.nlm.nih.gov/ReferenceSamples/giab/data/NA12878/analysis/PacBio_CCS_15kb_20kb_chemistry2_042021/ and here https://ftp-trace.ncbi.nlm.nih.gov/ReferenceSamples/giab/data/NA12878/HudsonAlpha_PacBio_CCS/.

### HG002 Cell Lines

The HG002 cell line R9 and R10 data is openly available through the AnVIL workspace: https://anvil.terra.bio/#workspaces/anvil-datastorage/ANVIL_NIA_CARD_Coriell_Cell_Lines_Open. An ultra-long HG002 data set was also used located here: https://s3.amazonaws.com/giab-aws/index.html?prefix=WGS/ONT/2022/11_16_22_R1041_HG002_UL_Kit14_400/.

## Code availability

https://github.com/NIH-CARD/CARDlongread_meth_R.9vs10

## Competing Interests

The authors declare no competing interests.

## Ethics Declarations

M.J. has received reimbursement for travel, accommodation and conference fees to speak at events organized by ONT.

## Acknowledgements

We would like to thank all of the participants who donated their time and biological samples to be a part of this study. This work utilized the computational resources of the NIH HPC Biowulf cluster (http://hpc.nih.gov). This work was supported in part by the Intramural Research Program of the National Institute on Aging (NIA) (AG000542-01, AG000538-03).This work was supported by the Center for Alzheimer’s and Related Dementias, within the Intramural Research Program of the National Institute on Aging and the National Institute of Neurological Disorders and Stroke, National Institutes of Health, Department of Health and Human Services (ZIAAG000538).

We thank members of the North American Brain Expression Consortium (NABEC, phs001300) for providing samples derived from brain tissue. Brain tissue for the NABEC cohort were obtained from the Baltimore Longitudinal Study on Aging at the Johns Hopkins School of Medicine, the NICHD Brain and Tissue Bank for Developmental Disorders at the University of Maryland, the Banner Sun Health Research Institute Brain and Body Donation Program, and from the University of Kentucky Alzheimer’s Disease Center Brain Bank.

Data biospecimens used in the analyses presented in this article were obtained from the Parkinson’s Progression Markers Initiative (PPMI) (www.ppmi-info.org/access-dataspecimens/download-data). As such, the investigators within PPMI contributed to the design and implementation of PPMI and/or provided data and collected biospecimens, but did not participate in the analysis or writing of this report. For up-to-date information on the study, visit www.ppmi-info.org. PPMI – a public-private partnership – is funded by The Michael J. Fox Foundation for Parkinson’s Research and funding partners, including 4D Pharma, AbbVie Inc., AcureX Therapeutics, Allergan, Amathus Therapeutics, Aligning Science Across Parkinson’s (ASAP), Avid Radiopharmaceuticals, Bial Biotech, Biogen, BioLegend, BlueRock Therapeutics, Bristol Myers Squibb, Calico Life Sciences LLC, Celgene Corporation, DaCapo Brainscience, Denali Therapeutics, The Edmond J. Safra Foundation, Eli Lilly and Company, Gain Therapeutics, GE Healthcare, GlaxoSmithKline, Golub Capital, Handl Therapeutics, Insitro, Janssen Pharmaceuticals, Lundbeck, Merck & Co., Inc., Meso Scale Diagnostics, LLC, Neurocrine Biosciences, Pfizer Inc., Piramal Imaging, Prevail Therapeutics, F. Hoffmann-La Roche Ltd and its affiliated company Genentech Inc., Sanofi Genzyme, Servier, Takeda Pharmaceutical Company, Teva Neuroscience, Inc., UCB, Vanqua Bio, Verily Life Sciences, Voyager Therapeutics, Inc., and Yumanity Therapeutics, Inc.

## Notes

### Competing Interest Statement

The authors have declared no competing interest.

