## Supplement - R9 and R10 5mC manuscript for "Assessing methylation detection for primary human tissue using Nanopore sequencing"

### Supplementary Materials

**Supplementary Figure 1:** Methylation intervals detected from R9 and R10 data for HG002 cell line **(a)**, brain sample **(b)**, and blood sample **(c)** binned according to R10 intervals. Distributions of CpG site frequencies for R9 and R10 are depicted on the right side of each of those subpanels. R9 methylation frequency distributions are shown in blue and the R10 distributions are shown in orange. The overlaid line plots connect the median interval points to visualize methylation trends.

a.

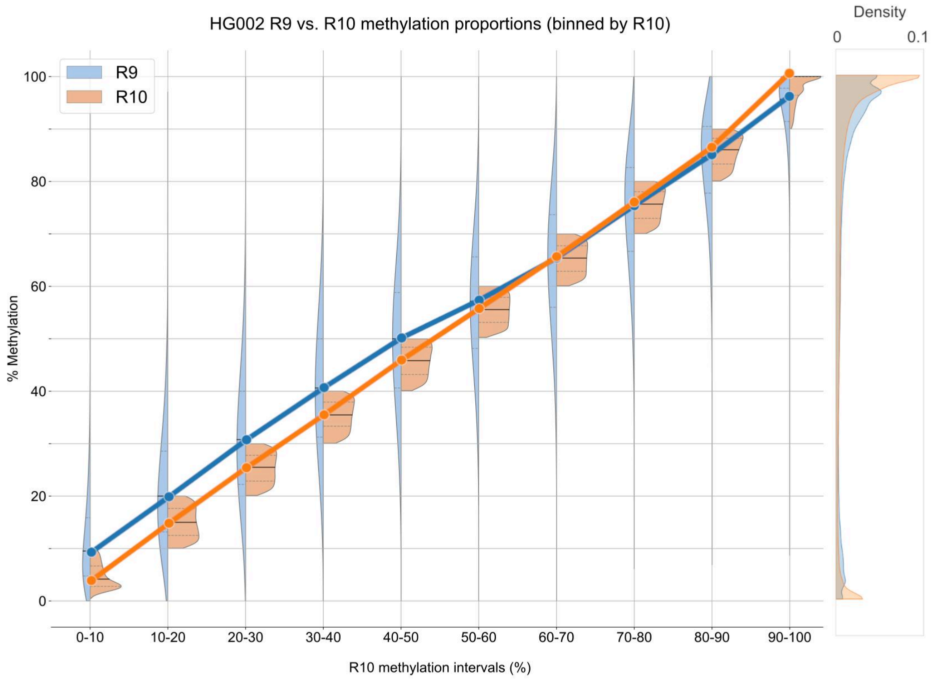


b.

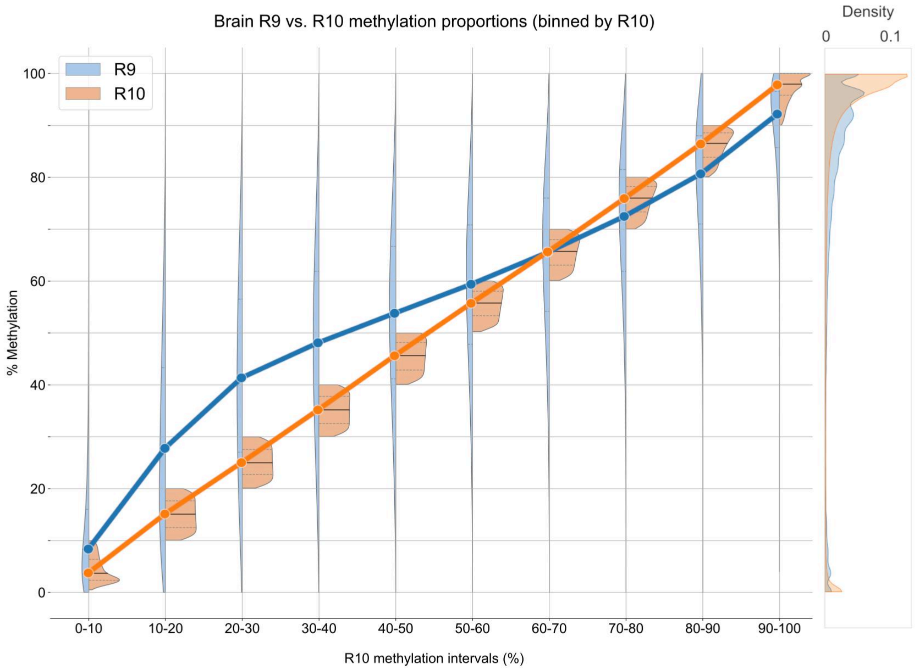
Brain R9 vs. R10 methylation proportions (binned by R10)
Density
0
0.1
R9
R10
100
80
60
40
20
0
% Methylation
0-10
10-20
20-30
30-40
40-50
50-60
60-70
70-80
80-90
90-100
R10 methylation intervals (%)

c.

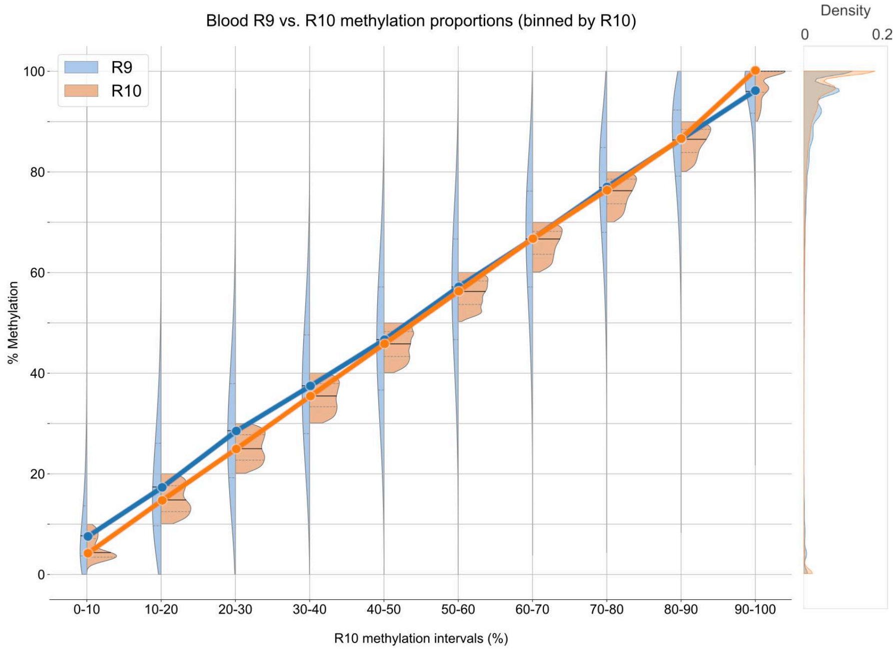
Blood R9 vs. R10 methylation proportions (binned by R10)
Density
0
0.2
R9
R10
100
80
60
40
20
0
% Methylation
0-10
10-20
20-30
30-40
40-50
50-60
60-70
70-80
80-90
90-100
R10 methylation intervals (%)

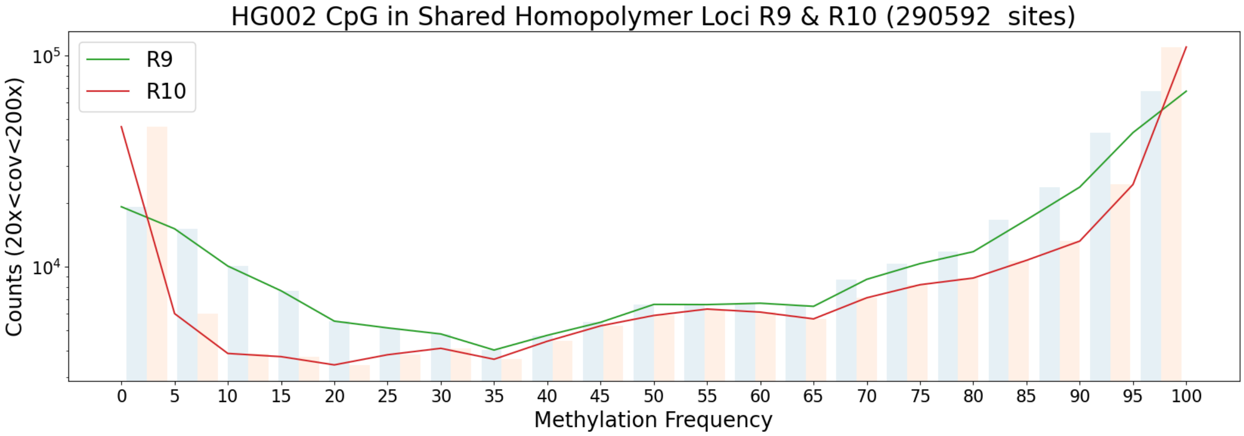


**Supplementary Figure 2.** Methylation frequency in shared homopolymer loci for R9 and R10.

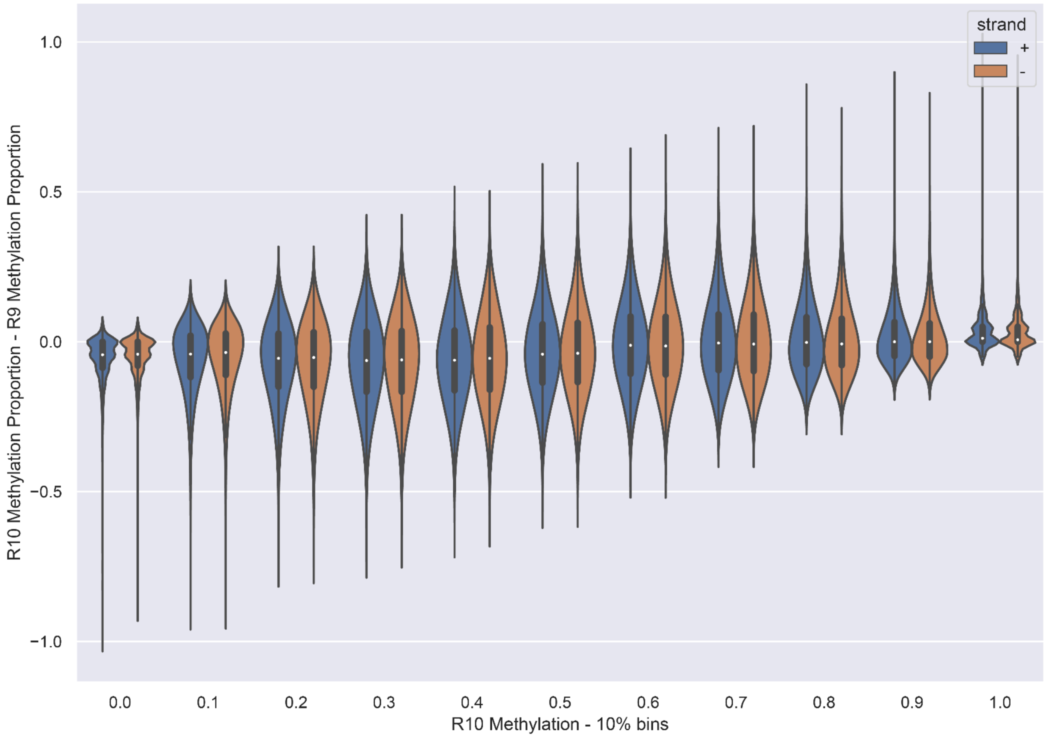


**Supplementary Figure 3.** Difference in methylation proportion between R10 and R9 datasets divided by strand of origin and binned by R10 methylation proportion.

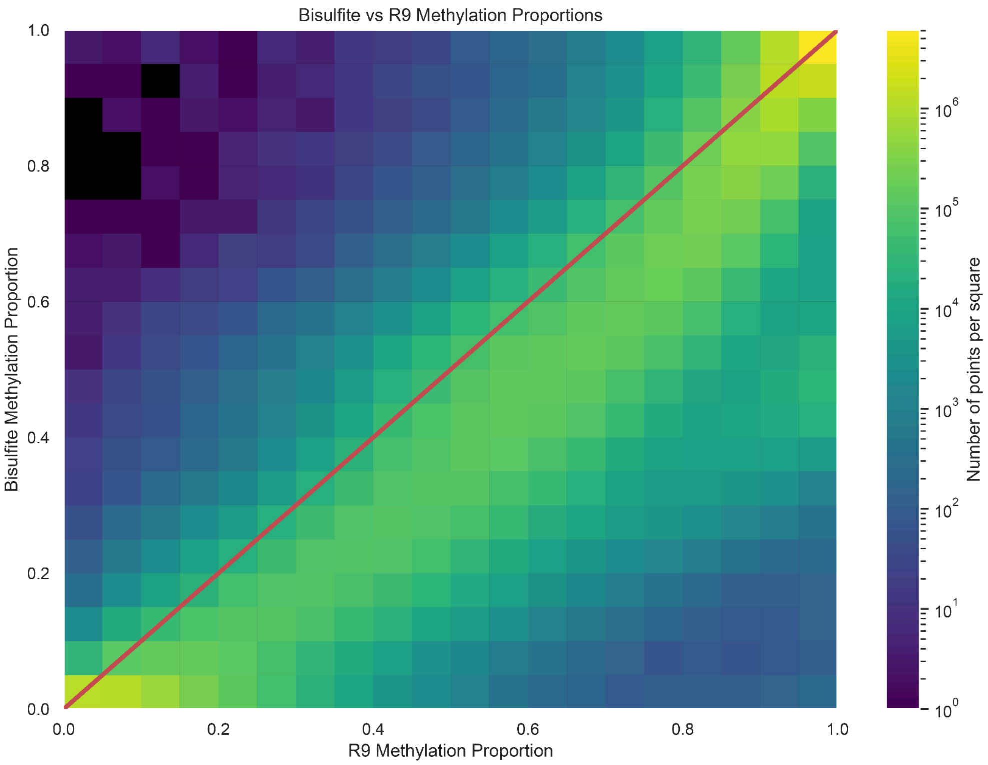


**Supplementary Figure 4.** R9 Ultra long methylation proportion data plotted against Bisulfite methylation proportion data.

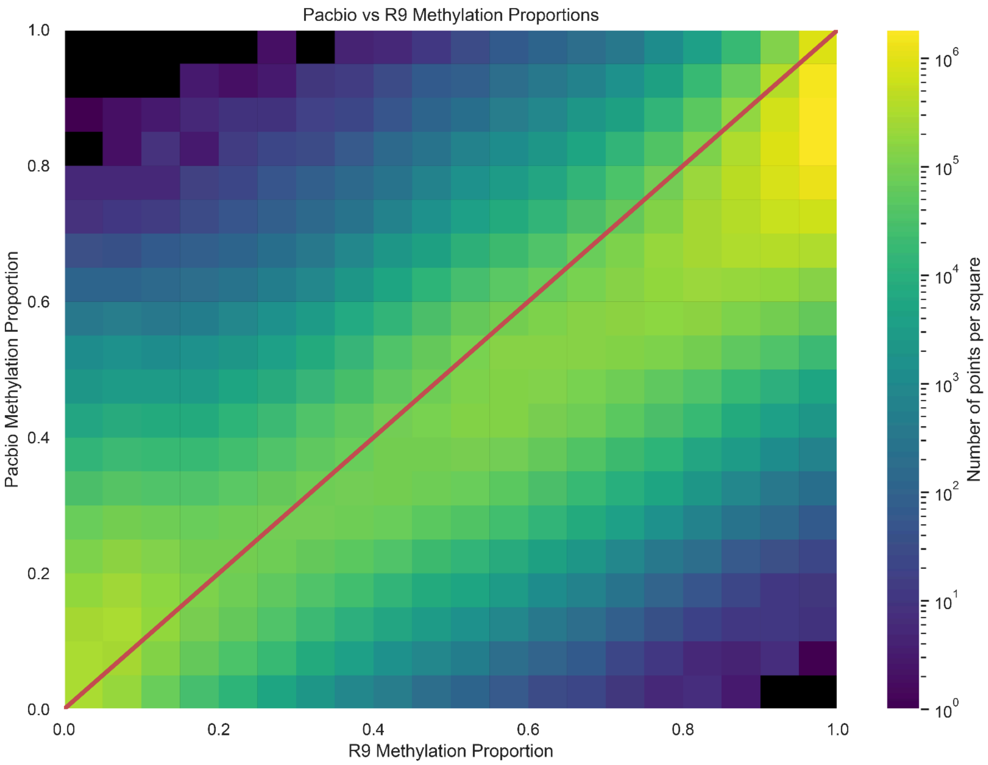


**Supplementary Figure 5.** R9 Ultra long methylation proportion data plotted against Pacbio methylation proportion data.

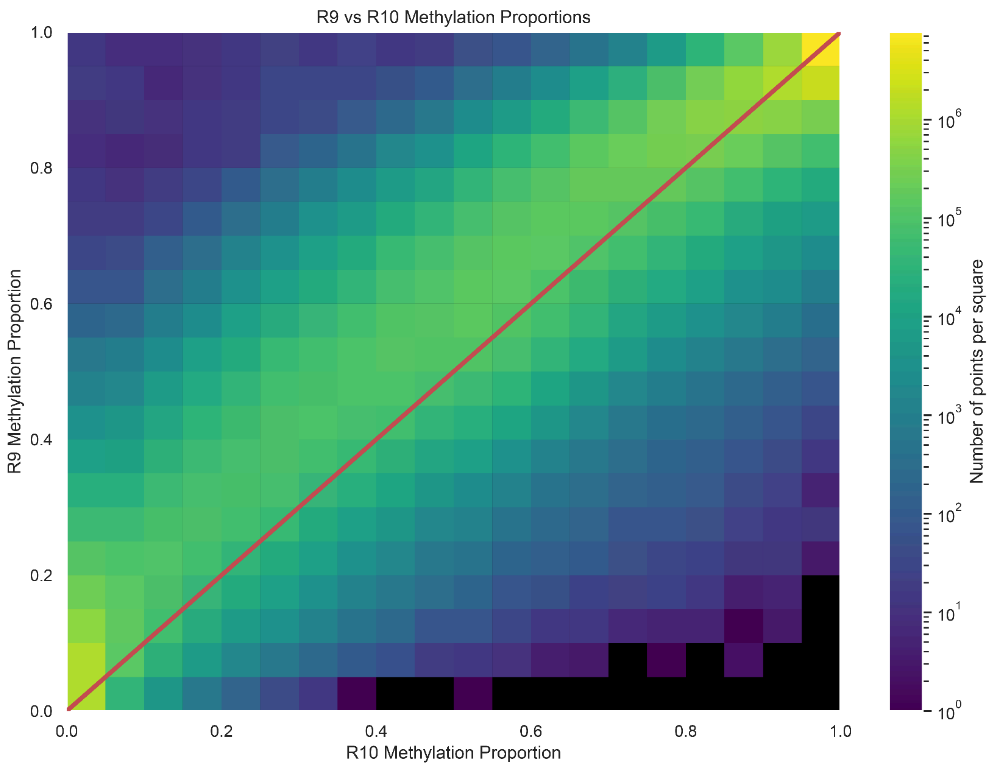


**Supplementary Figure 6.** R10 Ultra long methylation proportion data plotted against R9 Ultra long methylation proportion data.

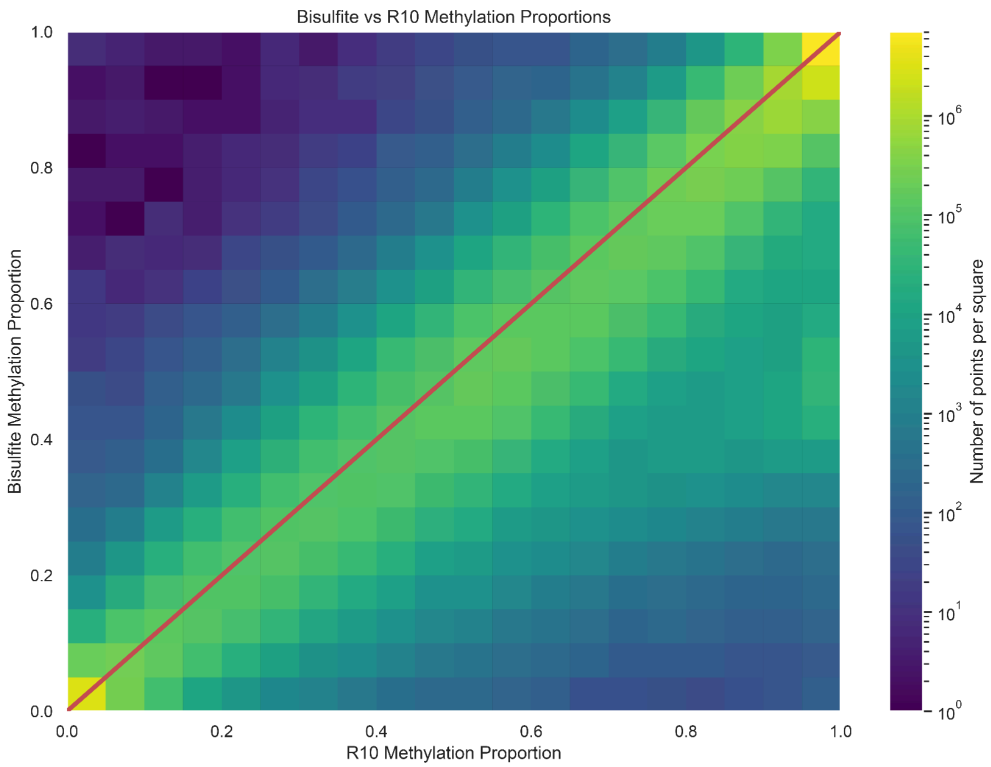


**Supplementary Figure 7.** R10 Ultra long methylation proportion data plotted against Bisulfite methylation proportion data.

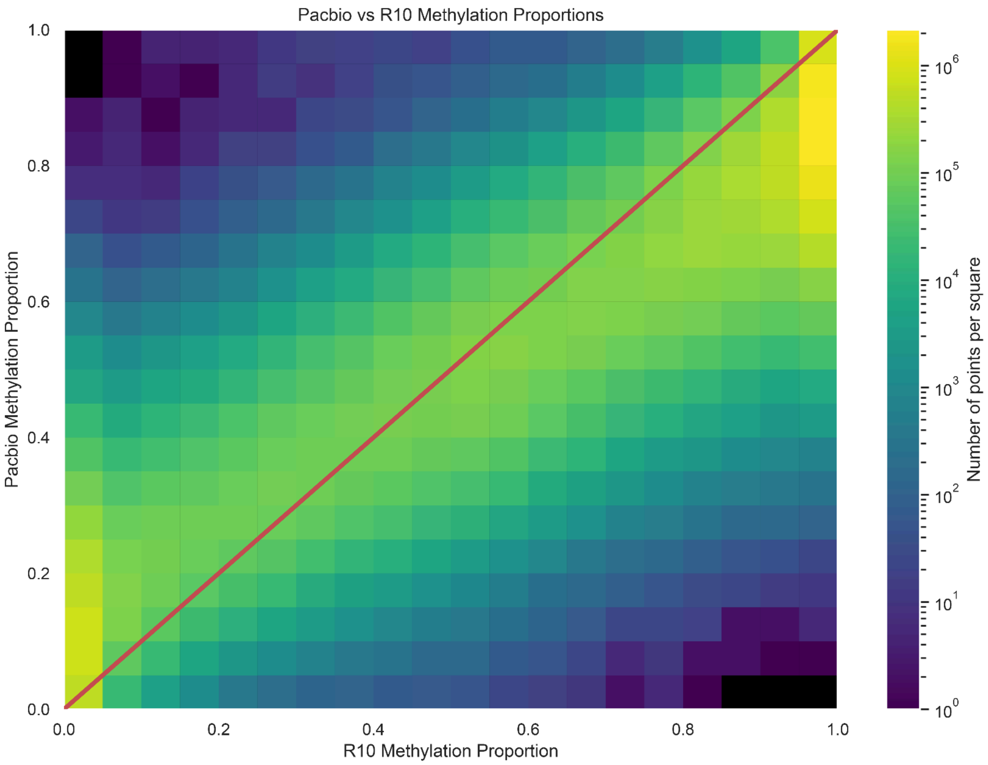


**Supplementary Figure 8.** R10 Ultra long methylation proportion data plotted against Pacbio methylation proportion data.

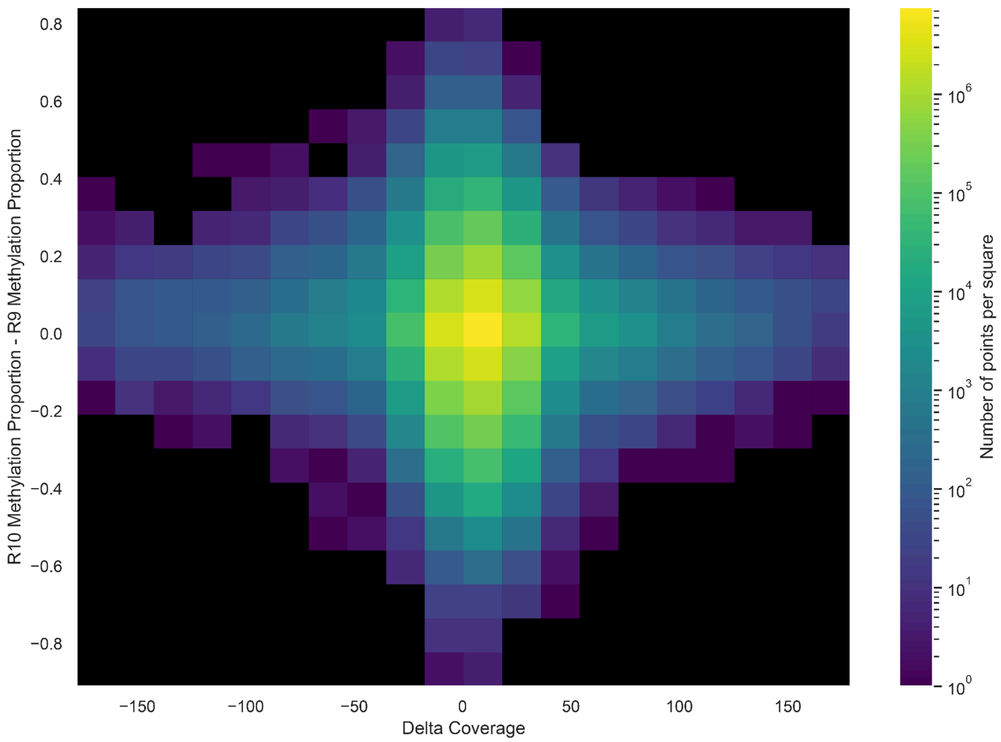


**Supplementary Figure 9.** Difference in coverage between R10 Methylation dataset and R9 Methylation dataset plotted against their difference in proportion.

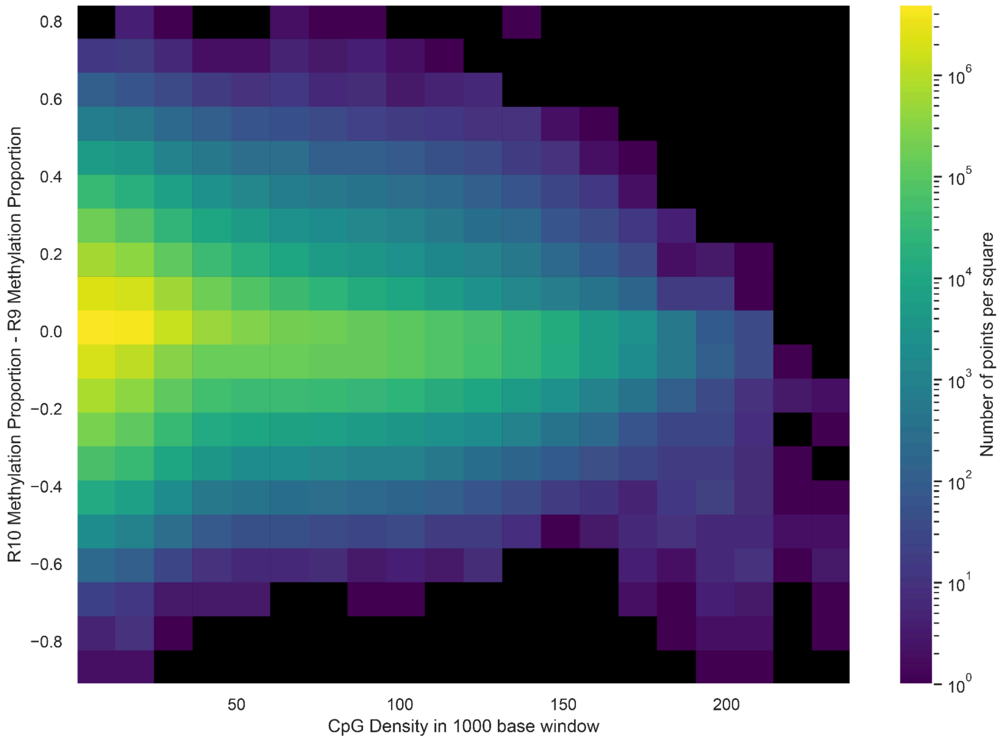


**Supplementary Figure 10.** Number of candidate CpG sites on the reference genome GRCh38 in a 1000 base window around the point of interest plotted against the difference in proportions of methylation of R10 ultra long and R9 ultra long data sets.

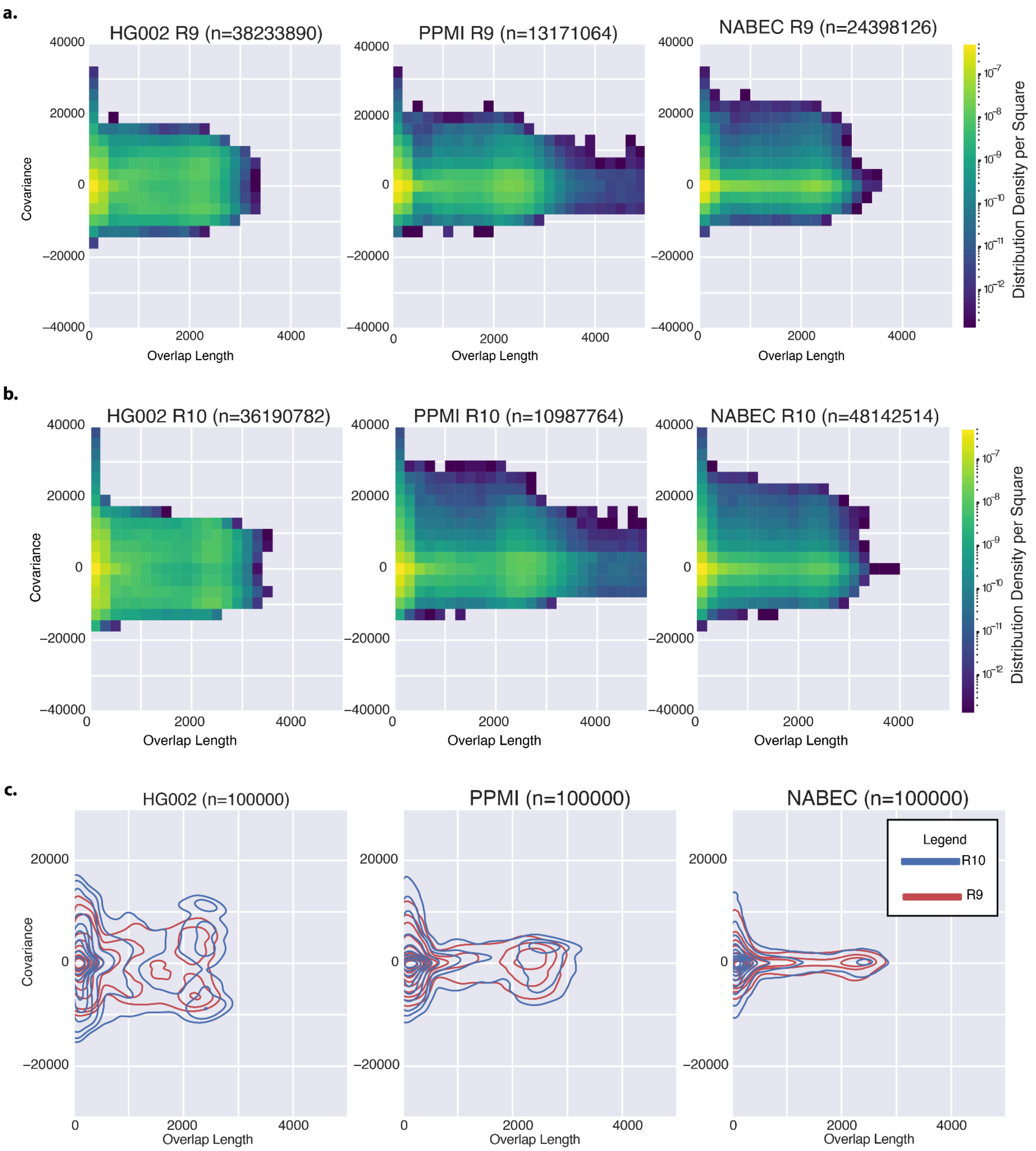


**Supplementary Figure 11:** a. Covariance measurements of modification confidence in overlapping sections (20+ CpG overlapping) of R9 reads in HG002, PPMI, and NABEC samples. b. Covariance measurements of modification confidence in overlapping sections of R10 reads in HG002, PPMI, and NABEC samples. c. Superimposed kernel density estimators of R9 and R10 covariance distributions for overlapping reads.

a.

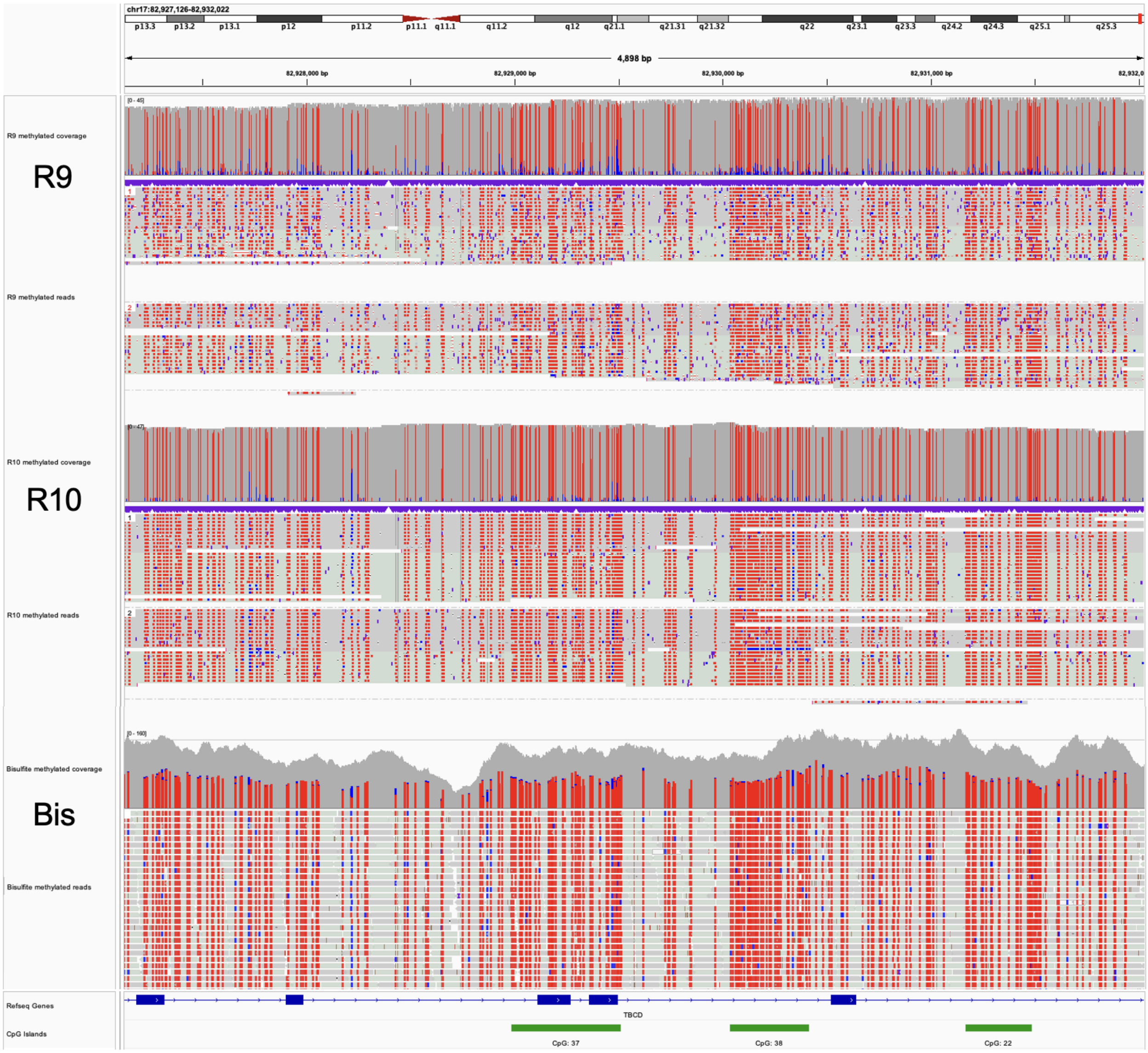

chr17:82,927,126-82,932,022
4,898 bp
R9 methylated coverage
R9
R9 methylated reads
R10 methylated coverage
R10
R10 methylated reads
Bisulfite methylated coverage
Bis
Bisulfite methylated reads
Refseq Genes
TBCD
CpG Islands
CpG: 37
CpG: 38
CpG: 22

b.

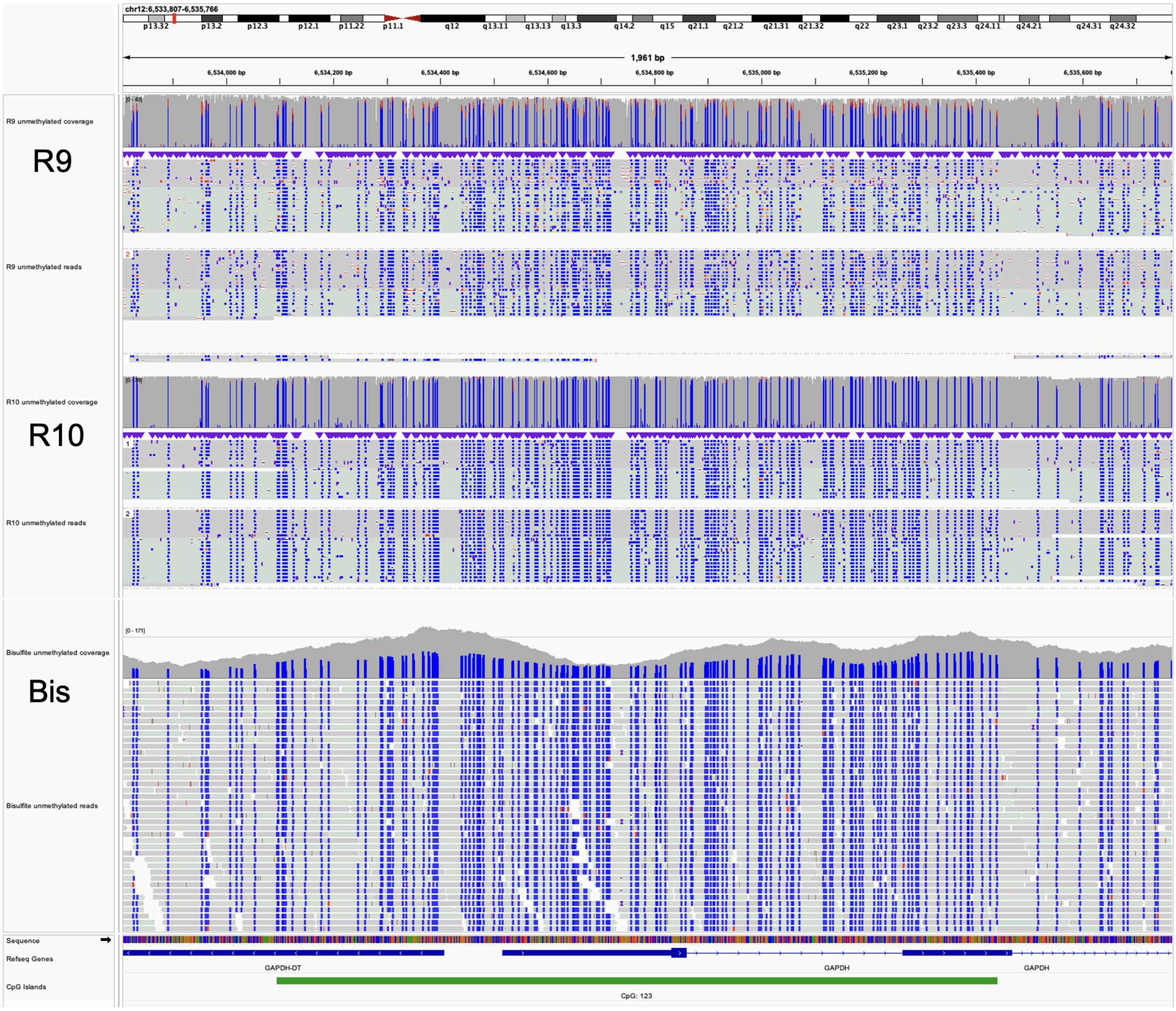


**Supplementary Figure 12. IGV plots depicting R9, R10, and bisulfite sequencing methylation differences in the constitutively methylated and unmethylated genomic regions in an HG002 cell line.** IGV [43] was used to visualize methylation patterns in HG002 cell line between ONT R9 and R10 methylation calls and traditional bisulfite sequencing in constitutively methylated (a) and unmethylated (b) regions. A CpG island associated with the GAPDH housekeeping gene was used as the constitutively unmethylated region (chr12:6533807-6535766)(a) and a region containing several “ultrastable” methylated CpG islands was used as the constitutively methylated region[41] (chr17:82927126-82923022)(b).

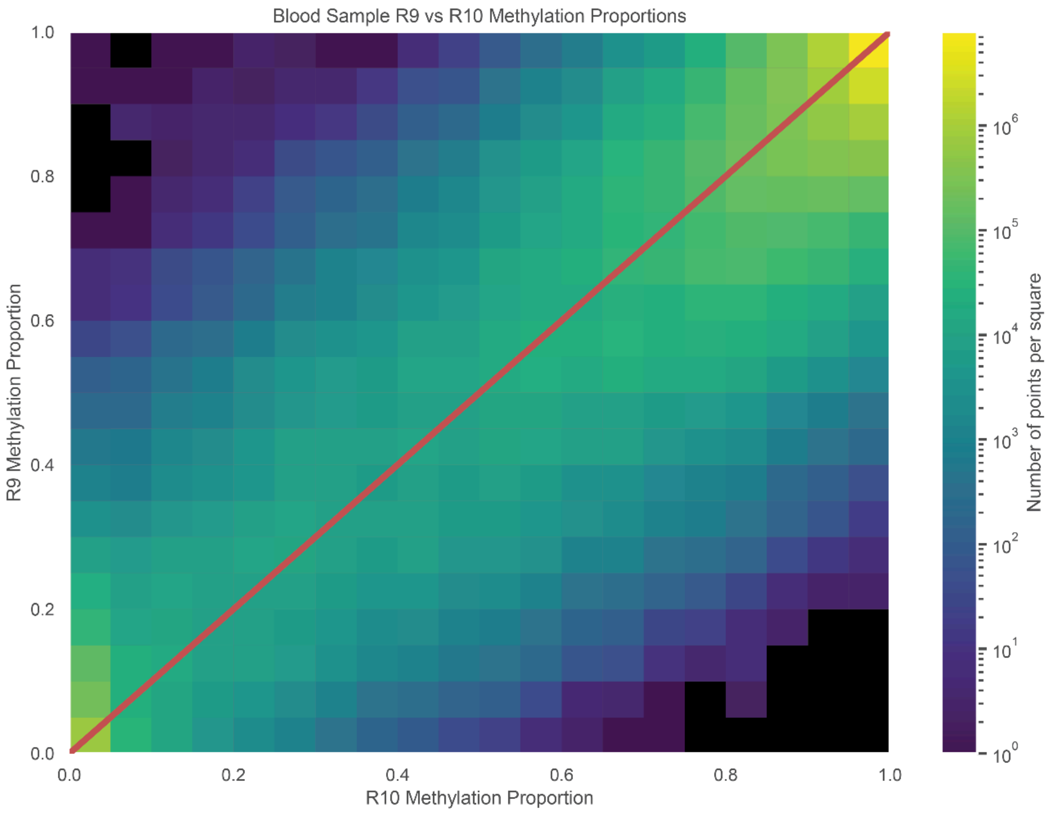


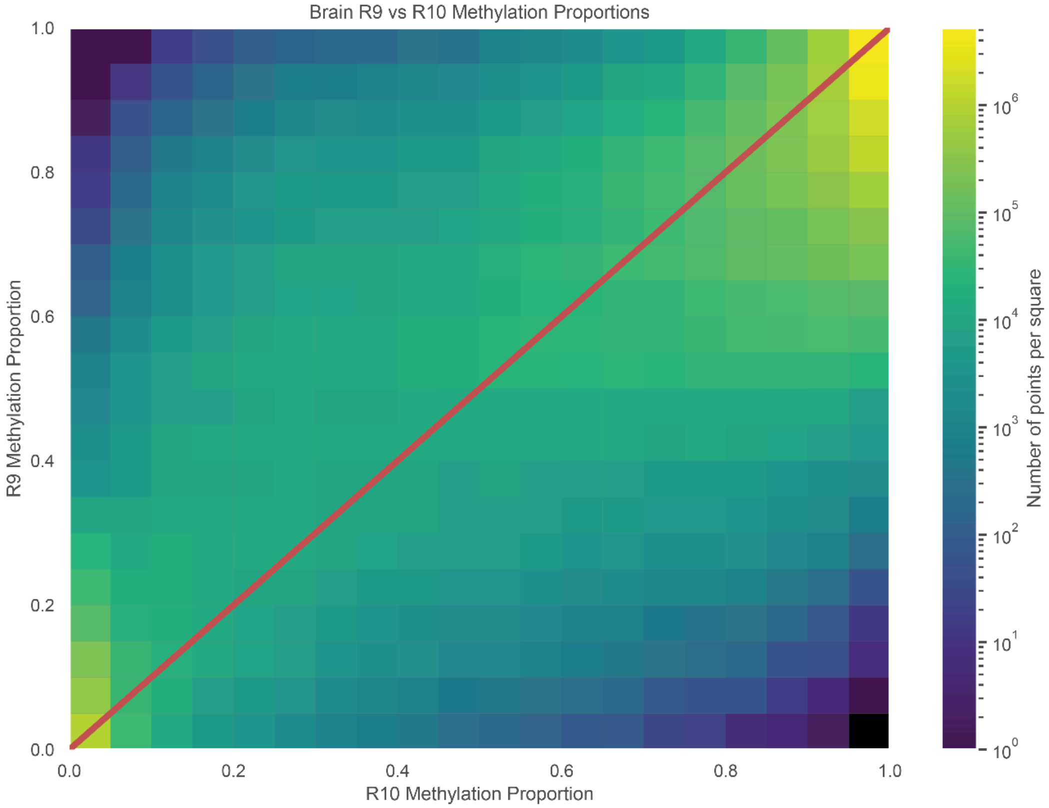


**Supplementary Figure 13.** top. Methylation proportion heatmap for Blood sample R9 and R10. bottom. Methylation proportion heatmap for Brain sample R9 and R10.

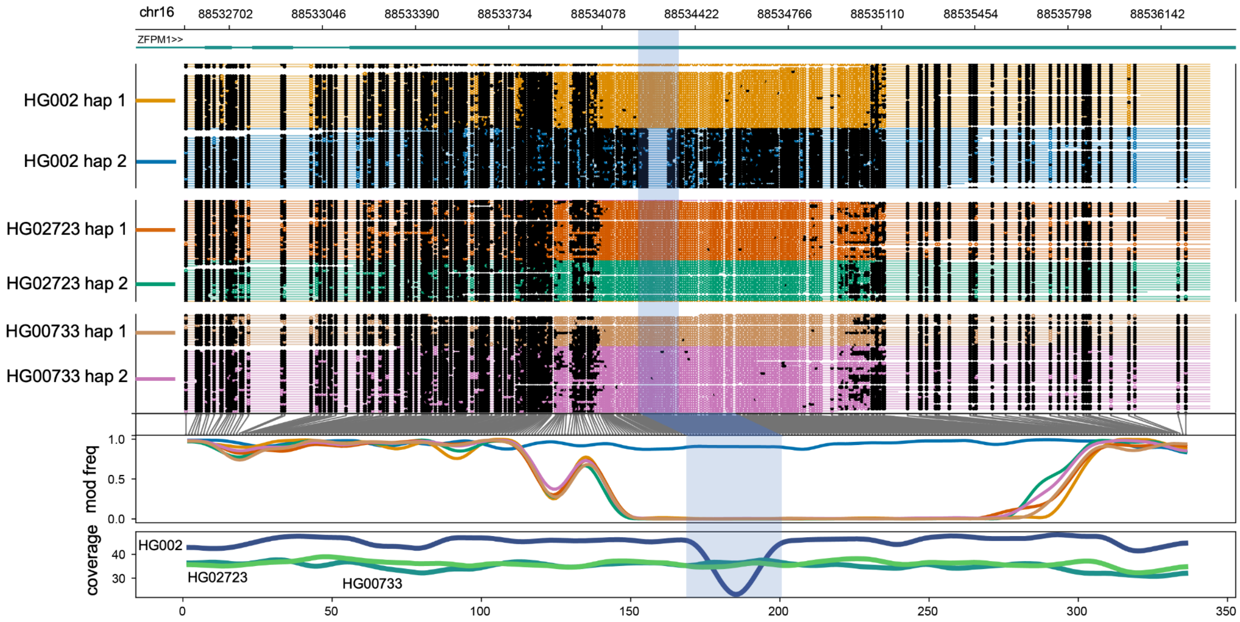


**Supplementary Figure 14.** Haplotype-specific methylation differences and similarities between R10 sequenced HG002, HG02723, and HG00733 GIAB cell line samples. The highlighted region corresponds to a 75 bp deletion present in haplotype 2 of the HG002 cell line that coincides with haplotype-specific methylation

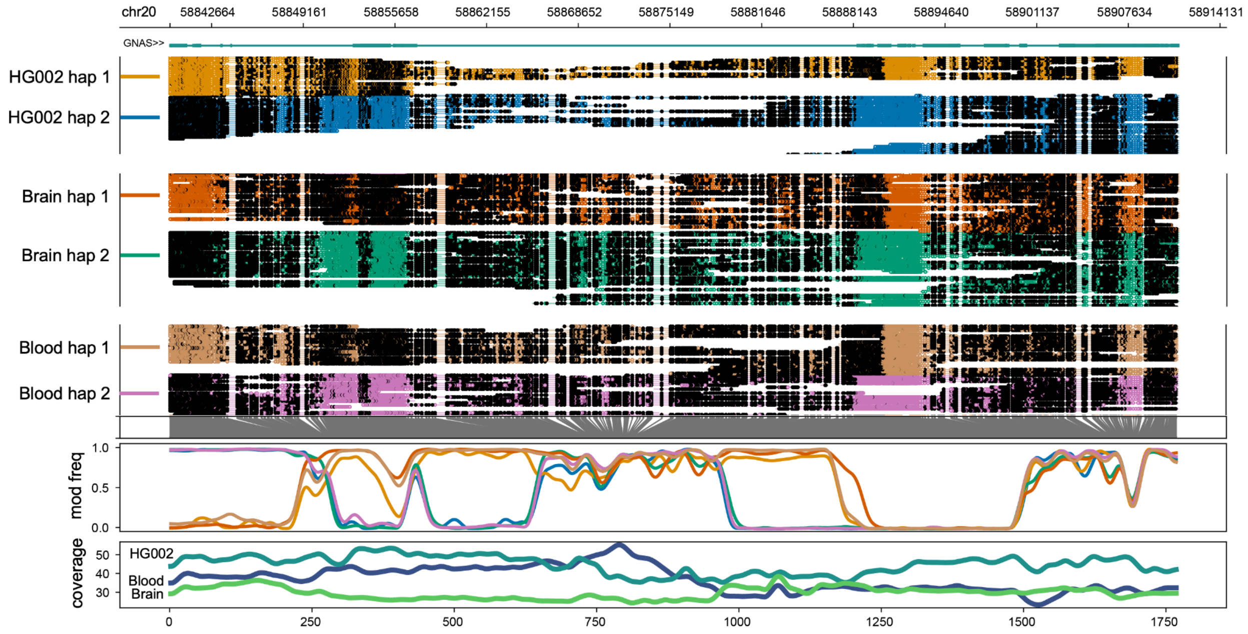


**Supplementary Figure 15.** Haplotype-specific methylation differences and similarities between the R10 sequenced cell, blood and brain samples in an imprinted region of the GNAS gene.

**Supplementary Table 1. Sequencing and alignment statistics for R9 and R10 across HG002, blood, and brain samples.**
All statistics were calculated using samtools md tags and pysam.

| | HG002 R9 | H002 R10 | Brain R9 | Brain R10 | Blood R9 | Blood R10 |
|---|---|---|---|---|---|---|
| Avg. cov | 42.1023 | 44.8986 | 38.8627 | 55.4873 | 38.8627 | 35.8224 |
| Avg. cov std | 127.921 | 113.94 | 96.7449 | 127.348 | 96.7449 | 91.704 |
| N50 (passed reads) | 27952 | 28483 | 30055 | 26026 | 34443 | 36484 |
| Alignment length mean | 17348.5724 | 14438.9332 | 20271.0092 | 7805.6603 | 24836.7853 | 18984.2836 |
| Alignment length median | 17129.0 | 10293.0 | 21580.0 | 1862.0 | 23912.0 | 14255.0 |
| Read identity mean | 0.9207 | 0.9657 | 0.9343 | 0.9693 | 0.9297 | 0.9601 |
| Read identity median | 0.9505 | 0.9872 | 0.952 | 0.9852 | 0.9536 | 0.9855 |
| Matches (per 1kb of ref bases) | 946.7014 | 978.5276 | 953.911 | 977.728 | 954.2411 | 975.7915 |
| Mismatches (per 1kb of ref bases) | 23.1096 | 9.6513 | 21.3427 | 9.9745 | 19.492 | 10.2314 |
| Deletions (per 1kb of ref bases) | 30.189 | 11.8212 | 24.7463 | 12.2975 | 26.2669 | 13.977 |
| Insertions (per 1kb of ref bases) | 17.0911 | 9.1746 | 16.5235 | 8.9238 | 15.8124 | 9.4844 |

**Supplementary Table 2. Comparing SNV calls between samples and technologies. SNV - Single Nucleotide Variant; INDEL - Insertion or Deletion; TS - Transitions; TV - Transversion.**

SNVs were for R9 and R10 HG002, brain, and blood samples were called using PEPPER-MARGIN-Deepvariant (PMDV) with R9 and R10 flags, respectively. SNV counts were calculated from the PMDV output vcf files for each sample using the bcftools stats package. Counts include: total single nucleotide variants (SNVs), insertions and deletions (INDELS), transitions (TS) and transversions (TV).

| | HG002 | | Brain | | Blood | |
|---|---|---|---|---|---|---|
| | R9 | R10 | R9 | R10 | R9 | R10 |
| SNVs | 4550624 | 5869049 | 5652676 | 5851663 | 4533415 | 5893333 |
| INDELS | 1353509 | 1220560 | 1314944 | 1198720 | 1358819 | 1247972 |
| TS | 2976460 | 3521071 | 4103611 | 3501067 | 2958780 | 3538987 |
| TV | 1577383 | 2363026 | 1552493 | 2365308 | 1578077 | 2368942 |

**Supplementary Table 3. Comparing SV calls between samples and technologies.**
SVs were called using sniffles2 and SV counts were calculated from the output vcf files for R9 and R10 sequenced HG002, brain, and blood samples. Counts include: total SVs present, deletions (DELS), insertions (INS), duplications (DUP), inversions (INV), and break-ends (BND).

| | HG002 | | Brain | | Blood | |
|---|---|---|---|---|---|---|
| | R9 | R10 | R9 | R10 | R9 | R10 |
| Total SVs | 26105 | 27428 | 26078 | 26778 | 26728 | 27643 |
| DELS | 11350 | 11905 | 11332 | 11678 | 11577 | 11916 |
| INS | 14627 | 15381 | 14631 | 14923 | 15047 | 15608 |
| DUP | 31 | 27 | 23 | 45 | 19 | 25 |
| INV | 48 | 46 | 43 | 52 | 48 | 44 |
| BND | 49 | 69 | 49 | 80 | 37 | 50 |

**Supplementary Table 4: CpG site capture for HG002, Blood, and Brain samples.**
Total CpG sites in bedMethyl file, filtered to >= 20 reads and <= 200 reads.

| | HG002 R9 | HG002 R10 | Blood R9 | Blood R10 | Brain R9 | Brain R10 |
|---|---|---|---|---|---|---|
| Total Sites | 28,798,028 | 28,760,288 | 28,623,063 | 28,606,058 | 28,814,883 | 28,798,897 |
| Filtered Sites | 25,937,319 | 27,021,032 | 22,347,084 | 25,723,371 | 23,271,407 | 27,742,379 |
| R9 and R10 Overlap | 25,521,492 | | 20,977,914 | | 23,148,718 | |

**Supplementary Table 5. Hyper and Hypo Methylated sites compared between chemistries (using R9 as a baseline) for HG002**

Identifying the number of sites in HG002 data where the extremes of methylation proportions are observed across chemistries (R9). Ratios refer to the methylation proportion at a given CpG site with >= 20 reads and <= 200 reads identified as canonical or modified.

| Proportion Conditions For HG002 | Site Count |
|---|---|
| R9 Ratio = 0 and R10 Ratio = 0 | 400,781 |
| R9 Ratio = 0 and R10 Ratio > 0 | 105,949 |
| 0 < R9 Ratio < 1 and 0 < R10 Ratio < 1 | 15,517,127 |
| 0 < R9 Ratio < 1 and R10 Ratio = 0 or R10 Ratio = 1 | 6,450,741 |
| R9 Ratio = 1 and R10 Ratio = 1 | 1,967,470 |
| R9 Ratio = 1 and R10 Ratio < 1 | 1,079,424 |

**Supplementary Table 6. Hyper and Hypo Methylated sites compared between chemistries (using R10 as a baseline) for HG002** Identifying the number of sites in HG002 data where the extremes of methylation proportions are observed across chemistries (R10). Ratios refer to the methylation proportion at a given CpG site with >= 20 reads and <= 200 reads identified as canonical or modified.

| Proportion Conditions For HG002 | Site Count |
|---|---|
| R10 Ratio = 0 and R9 Ratio = 0 | 400,781 |
| R10 Ratio = 0 and R9 Ratio > 0 | 1,826,337 |
| 0 < R10 Ratio < 1 and 0 < R9 Ratio < 1 | 15,517,127 |
| 0 < R10 Ratio < 1 and R9 Ratio = 0 or R9 Ratio = 1 | 1,185,373 |
| R10 Ratio = 1 and R9 Ratio = 1 | 1,967,470 |
| R10 Ratio = 1 and R9 Ratio < 1 | 4,624,404 |

**Supplementary Table 7. Hyper and Hypo Methylated sites compared between chemistries (using R9 as a baseline) for Blood Sample** Identifying the number of sites in Blood data where the extremes of methylation proportions are observed across chemistries (R9). Ratios refer to the methylation proportion at a given CpG site with >= 20 reads and <= 200 reads identified as canonical or modified.

| Proportion Conditions For Blood | Site Count |
|---|---|
| R9 Ratio = 0 and R10 Ratio = 0 | 368,132 |
| R9 Ratio = 0 and R10 Ratio > 0 | 52,108 |
| 0 < R9 Ratio < 1 and 0 < R10 Ratio < 1 | 10,432,127 |
| 0 < R9 Ratio < 1 and R10 Ratio = 0 or R10 Ratio = 1 | 5,041,706 |
| R9 Ratio = 1 and R10 Ratio = 1 | 3,257,111 |
| R9 Ratio = 1 and R10 Ratio < 1 | 1,826,730 |

**Supplementary Table 8. Hyper and Hypo Methylated sites compared between chemistries (using R10 as a baseline) for Blood Sample** Identifying the number of sites in Blood data where the extremes of methylation proportions are observed across chemistries (R10). Ratios refer to the methylation proportion at a given CpG site with >= 20 reads and <= 200 reads identified as canonical or modified.

| Proportion Conditions For Blood | Site Count |
|---|---|
| R10 Ratio = 0 and R9 Ratio = 0 | 368,132 |
| R10 Ratio = 0 and R9 Ratio > 0 | 630,146 |
| 0 < R10 Ratio < 1 and 0 < R9 Ratio < 1 | 10,432,127 |
| 0 < R10 Ratio < 1 and R9 Ratio = 0 or R9 Ratio = 1 | 1,878,838 |
| R10 Ratio = 1 and R9 Ratio = 1 | 3,257,111 |
| R10 Ratio = 1 and R9 Ratio < 1 | 4,411,560 |

**Supplementary Table 9. Hyper and Hypo Methylated sites compared between chemistries (using R9 as a baseline) for Brain Sample** Identifying the number of sites in Brain where the extremes of methylation proportions are observed across chemistries (R9). Ratios refer to the methylation proportion at a given CpG site with >= 20 reads and <= 200 reads identified as canonical or modified.

| Proportion Conditions For Brain | Site Count |
|---|---|
| R9 Ratio = 0 and R10 Ratio = 0 | 399,925 |
| R9 Ratio = 0 and R10 Ratio > 0 | 86,821 |
| 0 < R9 Ratio < 1 and 0 < R10 Ratio < 1 | 14,385,467 |
| 0 < R9 Ratio < 1 and R10 Ratio = 0 or R10 Ratio = 1 | 5,793,084 |
| R9 Ratio = 1 and R10 Ratio = 1 | 1,230,808 |
| R9 Ratio = 1 and R10 Ratio < 1 | 1,252,613 |

**Supplementary Table 10. Hyper and Hypo Methylated sites compared between chemistries (using R10 as a baseline) for Brain Sample** Identifying the number of sites in Brain data where the extremes of methylation proportions are observed across chemistries (R10). Ratios refer to the methylation proportion at a given CpG site with >= 20 reads and <= 200 reads identified as canonical or modified.

| Proportion Conditions For Brain | Site Count |
| --- | --- |
| R10 Ratio = 0 and R9 Ratio = 0 | 399,925 |
| R10 Ratio = 0 and R9 Ratio > 0 | 957,951 |
| 0 < R10 Ratio < 1 and 0 < R9 Ratio < 1 | 14,385,467 |
| 0 < R10 Ratio < 1 and R9 Ratio = 0 or R9 Ratio = 1 | 1,339,434 |
| R10 Ratio = 1 and R9 Ratio = 1 | 1,230,808 |
| R10 Ratio = 1 and R9 Ratio < 1 | 4,835,433 |

**Supplementary Table 11. Differentially methylated region computation statistics comparing haplotype-phased samples.**
Differentially methylated regions of HG002 R9 / R10 data were calculated with Nanomethphase DMA.

| | R9 Haplotype 1 vs. Haplotype 2 | R10 Haplotype 1 vs. Haplotype 2 |
|---|---|---|
| DMR Count | 10799 | 12782 |
| DMR CG Mean (SD) | 34.504 (44.487) | 33.485 (38.603) |
| DMR CG Count | 372612 | 428000 |
| Methylation Proportion Difference Mean (SD) | 0.2907 (0.1557) | 0.3143 (0.1859) |

**Supplementary Table 12. Differentially methylated region nucleotide overlap counts between chemistries.**
Differentially methylated regions compared between R9 and R10 chemistries.

| | R9 DMRs | R10 DMRs |
|---|---|---|
| Number of nucleotides shared between DMRs between chemistries | 5,125,887 | 5,125,887 |
| Number of nucleotides captured by DMRs | 6,739,192 | 8,216,370 |
